# Pan-cancer Analysis Reveals m^6^A Variation and Cell-specific Regulatory Network in Different Cancer Types

**DOI:** 10.1101/2023.12.11.571179

**Authors:** Yao Lin, Jingyi Li, Shuaiyi Liang, Yaxin Chen, Yueqi Li, Yixian Cun, Lei Tian, Yuanli Zhou, Yitong Chen, Jiemei Chu, Hubin Chen, Qiang Luo, Ruili Zheng, Gang Wang, Hao Liang, Ping Cui, Sanqi An

**Author notes:** Equal contribution (Yao Lin, Jingyi Li, Shuaiyi Liang, Yaxin Chen). **Corresponding authors:** (An S); (Cui P); (Liang H).

## Abstract

As the most abundant mRNA modification in mRNA, *N^6^*-methyladenosine (m^6^A) plays a crucial role in RNA fate, impacting cellular and physiological processes in various tumor types. However, our understanding of the function and role of the m^6^A methylome in tumor heterogeneity remains limited. Herein, we collected and analyzed m^6^A methylomes across nine human tissues from 97 m^6^A-seq and RNA-seq samples. Our findings demonstrate that m^6^A exhibits different heterogeneity in most tumor tissues compared to normal tissues, which contributes to the diverse clinical outcomes in different cancer types. We also found that the cancer type-specific m^6^A level regulated the expression of different cancer-related genes in distinct cancer types. Utilizing a novel and reliable method called “m^6^A-express”, we predicted m^6^A– regulated genes and revealed that cancer type-specific m^6^A-regulated genes contributed to the prognosis, tumor origin and infiltration level of immune cells in diverse patient populations. Furthermore, we identified cell-specific m^6^A regulators that regulate cancer-specific m^6^A and constructed a regulatory network. Experimental validation was performed, confirming that the cell-specific m^6^A regulator *CAPRIN1* controls the m^6^A level of *TP53*. Overall, our work reveals the clinical relevance of m^6^A in various tumor tissues and explains how such heterogeneity is established. These results further suggest the potential of m^6^A for cancer precision medicine for patients with different cancer types.

## Introduction

It is widely accepted that cancer is a disease caused by genomic and epigenetic changes in oncogenes and tumor suppressor genes. Pan-cancer analysis of whole genomes, enhancer expression, long noncoding RNA (lncRNA) regulation, immune response and the like, aiming to examine the differences and similarities across different tumor types, has received extensive attention [1–4]. As the most prevalent mRNA modification, m^6^A is reversibly regulated by various classical m^6^A regulators, including methyltransferases (*METTL3*, *METTL14*, *VIRMA*, *ZC3H13*, *WTAP*, *CBLL1*/*HAKAI* and *RBM15*/*RBM15B*) and m^6^A demethylases (*ALKBH5* and *FTO*), which mediate demethylation of different m^6^A sites [5–7]. m^6^A ‘readers’, *YTH* family proteins, and *IGF2BPs* recognize this reversible m^6^A and regulate post-transcriptional processes, such as RNA decay, alternative polyadenylation and nuclear export, in different cancer types [8, 9]. Previous studies have shown that m^6^A plays an important role in the progression of individual cancer types [10, 11]. However, our understanding of the mechanisms underlying cell and cancer type-specific m^6^A regulation in pan-cancer is limited [12, 13].

To investigate the complex cell type-specific regulatory network and the role of m^6^A in different cancer types, it is necessary to perform a pan-cancer analysis at the level of m^6^A methylation. Due to the limitations of m^6^A identification methods, some researchers have only tried to perform gene expression analysis of classical m^6^A regulators to analyze the heterogeneity of m^6^A regulators instead of real m^6^A in different cancers [14, 15]. However, key m^6^A regulators, such as *METTL3*, *METTL14* and *YTHDC2*, may function independently of m^6^A [16–18]. Therefore, these m^6^A regulators expression-based pan-cancer analysis, are far from revealing the characteristics and roles of m^6^A in different cancer types. Among the various methods for identifying m^6^A sites with high resolution, *N^6^*-methyladenosine sequencing (m^6^A– seq) is the most widely used and has promoted m^6^A research [19]. In our previous study, we made multiple methodological improvements to mitigate the impact of technical biases caused by different immunoprecipitation efficiencies across the different libraries in m^6^A-seq [13] to provide a reliable method for m^6^A pan-cancer analysis.

With accurate calculations based on a large number of m^6^A-seq datasets, we attempted to investigate the m^6^A landscape in terms of cancer tissue specificity. Then, we wanted to know what key clinically relevant heterogeneity of m^6^A leads to and how this heterogeneity of m^6^A is established. To achieve this, we performed comprehensive pan-cancer analyses involving nine cancer types utilizing 97 m^6^A-seq and RNA-seq samples. We used a reliable method called “m^6^A-express” to predict m^6^A-reg-exp genes (m^6^A regulation of gene expression) in pan-cancer analysis [12]. Through these m^6^A-reg-exp genes, we comparably and comprehensively explored clinically relevant m^6^A modifications that are involved in cancer progression, prognostic prediction and molecular classification across 31 different cancer types. Additionally, we attempted experimental validation of the *CAPRIN1*/*METTL3*-m^6^A– *TP53* axis identified by the in *silico* work. To the best of our knowledge, this is the first pan-cancer analysis based on global m^6^A levels. Our results will contribute to a better understanding of m^6^A heterogeneity and its role in cancer precision therapy

## Results

### Systematic analysis of extensive m^6^A-seq data reveals distinct m^6^A features in nine types of cancer and normal tissues

We collected m^6^A-seq data of 97 tumor tissues as well as corresponding normal tissues and explored the characteristics of these m^6^A sites. Various technical issues associated with m^6^A-seq data can obstruct the successful systematic analysis of quantitative m^6^A ratios calculated from m^6^A-seq data. As a result, we implemented multiple processing steps to minimize the effects of different types of technical issues (see the “Materials and Methods” section for details). Due to technical biases in preparing the m^6^A-seq libraries, such as variations in RNA length, the shifting of peak centers and divergence of peak widths will be controlled using winscore methods (see the “Materials and Methods” section for details). Afterwards, quantile normalization of the m^6^A ratios is performed to correct biases introduced by differences in antibody efficiencies across various laboratories. Based on our previous findings that m^6^A sites with a CV greater than 0.3 vary between different cells [13], we further calculated the specificity and function of these m^6^A sites. Subsequently, we utilized the expression levels of cell-specific m^6^A regulators identified in our previous research and classical m^6^A regulators to compute their correlation with the m^6^A we obtained. Through experimental validation, we identified specific regulators that modulate cancer-specific m^6^A. At the same time, we employed m^6^A-express to calculate target genes regulated by m^6^A in different cancers and analyzed the potential impact of these m^6^A– regulated genes on prognosis and cellular immune responses across various cancers (Figure 1).

**Figure 1.**
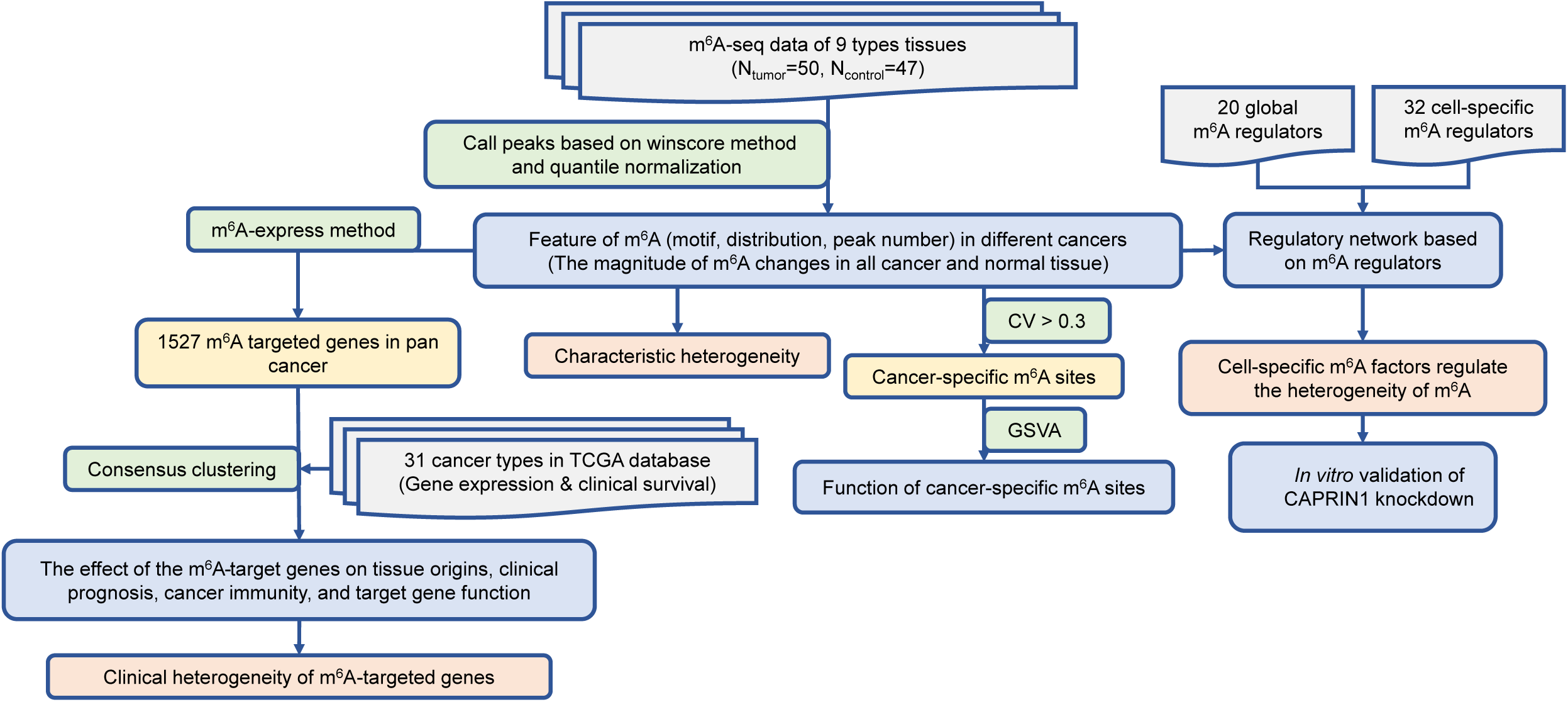
Schematic fow chart demonstrating the process of the analysis.

To systematically reveal the basic characteristics of m^6^A in various cancer types at the tissue level, we initially made several methodological improvements to the winscore-based method to mitigate the impacts of technical biases of m^6^A-seq data from different laboratories (see the ‘Methods’ section for more details) [13]. This is important for the reliable quantification of m^6^A levels using m^6^A-seq data from different labs. After performing quantile normalization of the m^6^A level across all tissues, we performed unsupervised clustering using the normalized m^6^A level and found that the variable m^6^A levels in lung cancer and leukemia were not clustered according to the labs but rather according to cancer types, suggesting that we successfully eliminated batch effects from different labs (Figure S1A). Then, we conducted m^6^A quantitative analysis on nine tumor tissues and their controls, including glioma, lung cancer, liver cancer, endometrial cancer, ovarian cancer, leukemia, colon cancer, salivary gland cancer, and stomach cancer. Approximately 10,000 m^6^A peaks were obtained for each tissue (Figure 2A), and samples from the same tissue type clustered well according to their m^6^A levels (Figure 2B), indicating that *tissue-specific* m^6^A methylomes indeed exist. To better understand the biological function of these m^6^A sites, we annotated them and found that m^6^A mostly occurred on protein-encoding genes (Figure S1D). Some m^6^A was present on non-coding RNA (Figure S1D), suggesting that non-coding RNA also played essential roles in these cancer types. We observed a trend that m^6^A from different cancer types had varying levels of enrichment in distinct m^6^A sub-motifs (*P* < 2.2×10^-16^), suggesting that m^6^A methylomes may be cancer-specific (Figure 2C, Figure S1E). Cancer-specific m^6^A motifs, m^6^A distribution, and m^6^A peak number indicated that m^6^A with different functions in different tumors is regulated in a cancer-specific manner.

**Figure 2.**
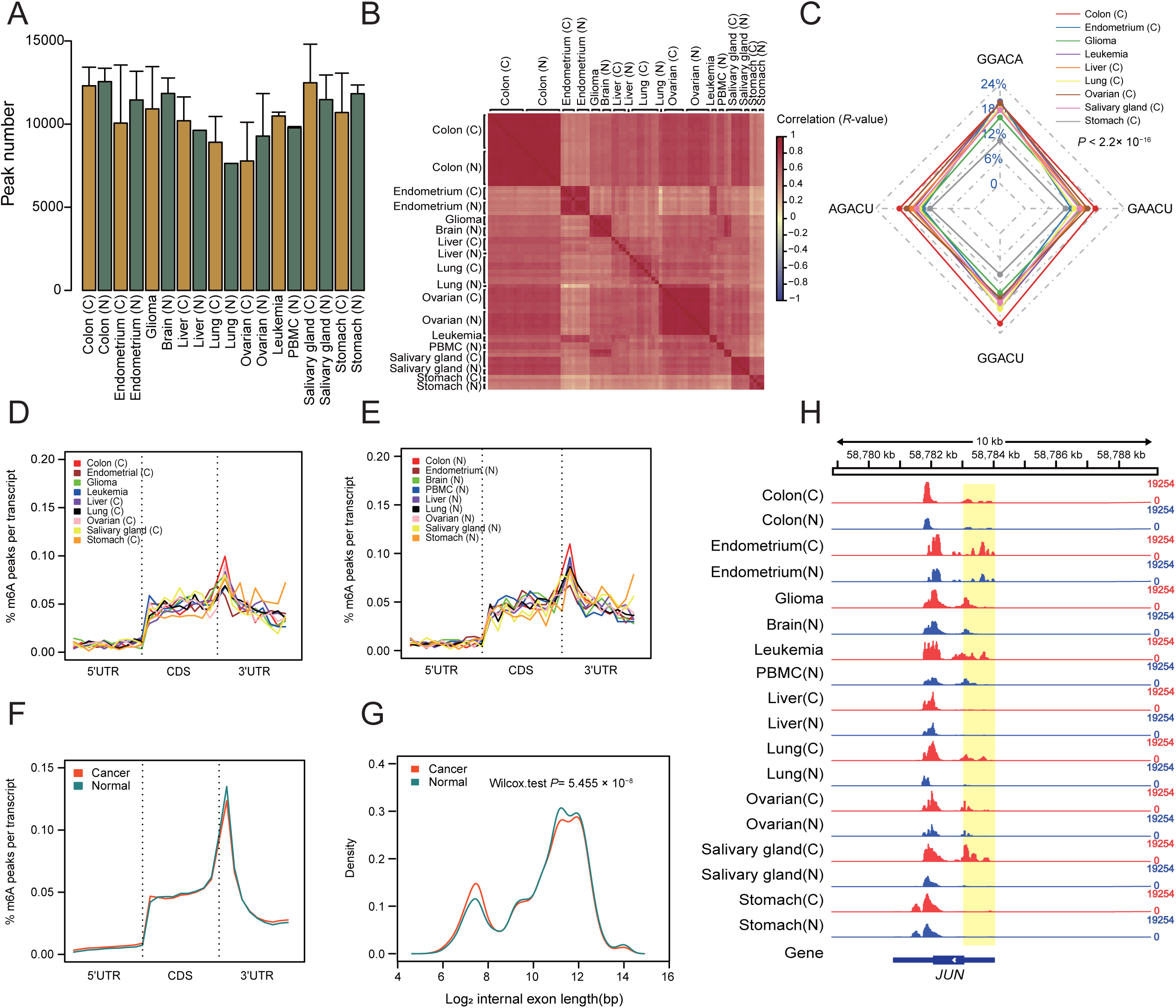
m^6^A features in nine tumor and normal tissues. **A**. The number of m^6^A peaks identified in each cancer and normal tissue is represented by a bar graph. Error bars denote the maximum and minimum across all replicates. **B.** Pearson correlation heatmaps of the correlation matrix representing the correlation between global m^6^A levels in different samples. **C.** The radar map showing the percentage of m^6^A sub-motifs of nine cancer tissues, indicated by different colored lines (Chi-Squared Test, *P* < 2.2×10^-16^). Normalized distributions of the m^6^A peak at 5’UTR, CDS, and 3’UTR in nine tumor tissues (**D**) and nine normal tissues (**E**). **F.** Comparison of overall m^6^A peak distribution between tumor and normal tissues. **G.** Exon length of tumor tissues compared with normal tissues. The *P* value of the Wilcoxon test is indicated (*P* = 5.455×10^-8^). **H.** Track showing m^6^A coverage of the gene *JUN* from randomly selected samples among 97 cancer and normal subjects, with the 5 ‘UTR highlighted. The data range for each track is displayed on the left side (0-19254).

Then, we studied the distribution of m^6^A in different samples (Figure 2D-E), and consistent with other reports, m^6^A was more likely to be enriched in the 3’UTR start segment and the CDS segment and reached its peak at the 3’UTR start segment. By integrating the distributions of tumor samples and control samples, it was also found that m^6^A peaks in tumor samples are more prevalent at stop codon regions than in normal samples, whereas m^6^A peaks in normal samples were more abundant at start codon regions (Figure 2F) and less abundant in long internal exons (*P*= 5.455 × 10^−8^) (Figure 2G). In our previous research, we considered the m^6^A sites away from stop codons as “dynamic m^6^A sites”. They are precisely and dynamically regulated by cell-specific *trans* regulators expressed with spatial and temporal specificities across different cell types [13]. Herein, we suspect that m^6^A in cancer tissues had a greater coefficient of variation (CV) than that of normal tissue, which might be regulated by cell-specific *trans* regulators and contribute to the heterogeneity of cancer. We also used *JUN*, which has been associated with human cancer malignancies, as an example to analyze the differences in the CV of m^6^A between cancer and normal tissues. At the beginning of the *JUN* gene coding region, higher CV on the start codons were observed in nine types of cancer tissues (Figure 2H).

To further validate our conclusions, we also gathered m^6^A-seq data from cancer and normal cell lines for further analysis, including 7 tumor cell lines (HEC1a, HEPG2, iSLK, MOLM13, MonoMac6, MT4tCELL, NB4) and 3 normal cell lines (MSC, NHDF, TIME) (Table S2). By analyzing the distribution and motif of m^6^A site characteristics, we found that m^6^A in normal cell lines tended to be closer to stop codons, whereas there were more m^6^A peaks in cancer cell lines in coding regions (Figure S1B), and there were differences in the enrichment of m^6^A motifs in different tumor types (Figure S1C). Overall, the analysis results of m^6^A-seq data from cell lines are consistent with those from tumor tissues, further indicating the reliability of our findings in tumor tissue.

### m^6^A shows different heterogeneity in most tumor tissues compared with normal human tissues

The CVs of m^6^A were calculated in different regions of mRNA to explore whether the CVs in different cancer samples were greater than those in normal tissues. There were 15–50% variable peaks in normal tissues (Figure S2A), consistent with a recent report that *cis* regulation accounts for 33–46% of the variability in m^6^A levels [20]. Moreover, the m^6^A peaks located far from stop codons had significantly higher CVs in colon cancer, endometrial cancer, lung cancer, salivary gland cancer and stomach cancer (Figure 3A–D, Figure S2B). Therefore, there were more variable peaks in these cancer samples than in normal samples, suggesting that m^6^A in these five cancer types tends to be variable. We also calculated the m^6^A CV fold-change of cancer with respect to normal tissues in start and stop codons. Figure 3E shows that m^6^A enriched in start codons tends to have higher CVs in cancer tissues than m^6^A in stop codons. Moreover, we used the *TP53* and *HSPD1* genes as examples to study variations in m^6^A in cancer, and found that *TP53* and *HSPD1* showed higher variations in m^6^A at the start of coding sequences in colon cancer and endometrial cancer than in normal tissues (Figure 3F, Figure S2C).

**Figure 3.**
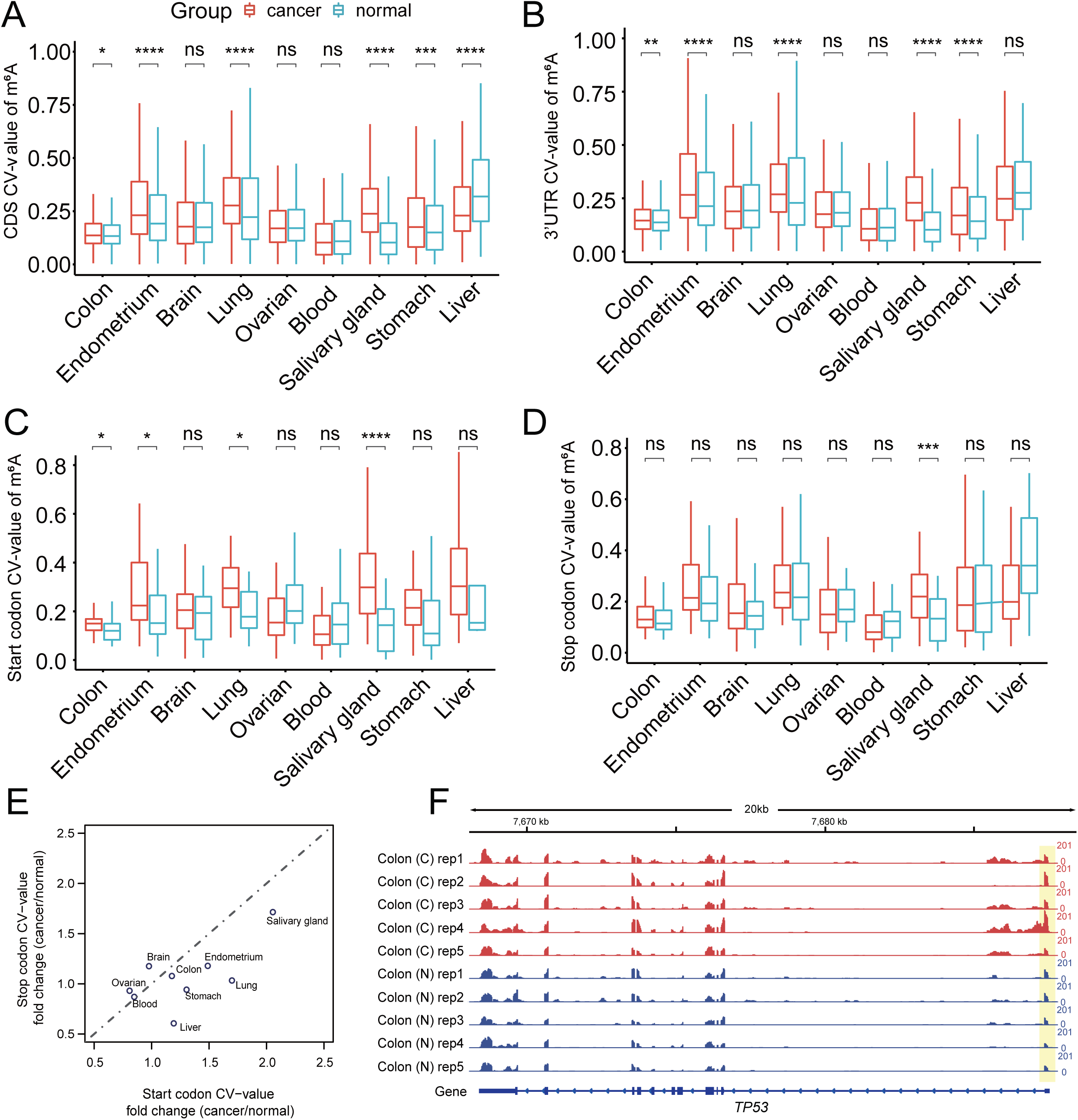
Coefficient of variation (CV) of the m^6^A level in tumor and normal tissues. Boxplot showing CV of the m^6^A level at the 3’UTR (**A**), CDS (**B**), and start codon (**C**), and stop codon (**D**) segments (*, *P* < 0.05; **, *P* < 0.01; ***, *P* < 0.001; ****, *P* < 0.0001). **E.** Dot plots representing fold change (cancer/normal) of coefficient of variation (CV) of m^6^A in the start and stop codons across nine tissue samples. **F.** The track displays the m^6^A abundance of the gene *TP53* in a lung tumor, with the 5’UTR highlighted. The data range for each track is displayed on the left side (0-201).

In addition to heterogeneity, we explored whether m^6^A in different cancer types had homogeneity. The samples were divided into two groups: the cancer group (45 samples) and the normal group (39 samples). We performed whole-genome different m^6^A analysis between these two groups. Only 428 m^6^A sites were differentially methylated [false discovery rate (FDR) < 0.05] between pan-cancer and normal tissues. m^6^A at these sites showed higher heterogeneity (*P* < 2.2 × 10^-16^) (Figure S2D). Thus, our findings indicate that m^6^A displays different heterogeneity in different tumor tissues.

### Tumor type-specific m^6^A affects distinct cancer-related genes and immune-related gene expression

To delve deeper into the role of cancer-specific m^6^A across various cancers, we identified m^6^A sites with a CV (coefficient of variation) greater than 0.3 as cancer-specific in tumor samples. This is because we observed that m^6^A sites with a CV greater than 0.3 often exhibit cell-specific characteristics, as previously noted [13]. Clustering showed that m^6^A levels were strongly cancer type-specific (Figure 4A). We also performed gene expression analysis of m^6^A-corresponding genes across all nine tumor tissues and found that gene expression may correlate with gene expression (Figure 4A-B). To further investigate the relationship between m^6^A and gene expression, we analyzed the correlation between the level of m^6^A modification at each site and the corresponding gene expression (Figure 4C). We observed that the majority of m^6^A modifications exhibited a positive correlation with the corresponding gene expression. This trend significantly differs from the results of randomly shuffled data for this correlation (*P* < 2.2 × 10^-16^). This outcome indicates that m^6^A modifications may be positively associated with gene regulation in tumor samples. This conclusion provides a valuable reference for guiding subsequent experimental research.

**Figure 4.**
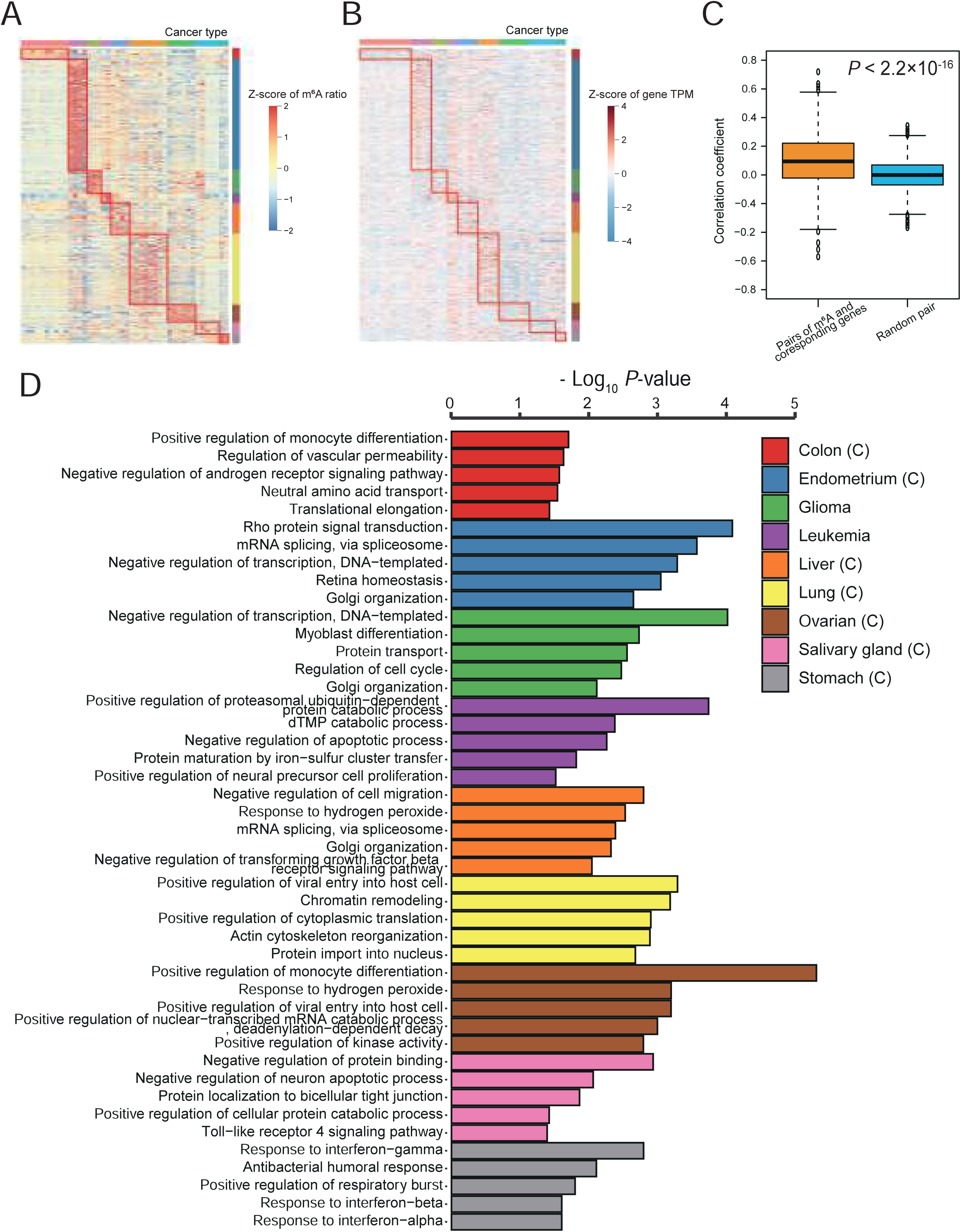
Relationship and functions between cancer type-specific m^6^A level and cancer type-specific gene expression. **A**. The m^6^A level in nine cancer types is displayed by heatmap. Each red box represents the cancer type-specific m^6^A site of each cancer. **B.** Heatmap showing the expression level of m^6^A targeted gene after log_2_ conversion to TPM in nine cancer types. Each red box represents the cancer type-specific m^6^A targeted gene of each cancer. **C.** The correlation between the level of m^6^A modification in each profile and the corresponding gene expression. The trend significantly differs from the results of randomly shuffled data for this correlation (*P* < 2.2×10^-16^). **D.** GO analysis of cancer-specific m^6^A-targeted genes in each cancer type.

GSVA enrichment analysis was employed to explore the biological functions among these distinct m^6^A modification patterns, and the results showed that different cancer-specific m^6^A was involved in different regulatory pathways, including various immune-related pathways, highlighting the relevance of m^6^A and tumor immunity (Figure S3). For example, colon cancer-specific m^6^A was enriched in pathways such as natural killer cell mediated cytotoxicity, cytokine receptor interaction, Jak stat signaling pathway, and Notch signaling pathway. Endometrial cancer-specific m^6^A was involved in B-cell receptor signaling pathway, T-cell receptor signaling pathway, and chemokine signaling pathway. Salivary gland cancer-specific m^6^A was associated with epithelial cell signaling in *Helicobacter pylori* infection and Toll-like receptor signaling pathway. Ovarian cancer-specific m^6^A was involved in autoimmune thyroid disease. In addition, in some cases, cancer-specific m^6^A was *tissue-specific*. For example, glioma-specific m^6^A was essential in neurotrophin signaling pathway and TGF beta signaling pathway. Leukemia-specific m^6^A was related to acute myeloid leukemia and cell cycle pathway. We also identified cancer-specific m^6^A that was significantly involved in the regulation of key tumor pathways. For instance, liver cancer-specific m^6^A was related to the WNT signaling pathway, lung cancer-specific m^6^A was enriched in glycerophospholipid metabolism, and stomach cancer-related m^6^A was involved in oxidative phosphorylation. These results illustrate the wide and varied regulatory mechanisms of m^6^A in different tumors.

We also performed Gene Ontology (GO) enrichment analysis (Figure 4D, Figure S4A and S4B). The analysis revealed significant enrichment of m^6^A in metabolic pathways, such as pyrimidine and purine and drug metabolism, as well as immune related pathways, including MAPK signaling, T-cell and B-cell pathways, leukocyte migration, infection, and inflammation. Analysis of cellular components showed that in pan-cancer, m^6^A was involved in the cell nucleus, cell membrane, Golgi apparatus, mitochondria, cytoplasm, endoplasmic reticulum, and exosome. These results are consistent with previous reports that m^6^A regulates tumor immunity and key tumor pathways [10]. Furthermore, these findings revealed that cancer-specific m^6^A tended to be enriched in different pathways and functions, highlighting the complex regulatory and functional specificity of m^6^A in different tumor types.

### The classification of different patients based on m^6^A-regulated genes was correlated with microenvironment and tissue origin

To explore the impact of m^6^A on tumorigenesis and development in pan-cancer, m^6^A– express, the first well-established algorithm to predict condition-specific m^6^A regulation of gene expression from MeRIP-seq data [12], was used to screen m^6^A-regulated genes in different cancers. A total of 1527 m^6^A-reg-exp genes were found in this article (Table S3).

These m^6^A-regulated genes were analyzed in 9456 tumor samples from 31 cancers for cluster analysis (Figure S5A and S5B), and it was found that pan-cancer related m^6^A-reg-exp genes could be used for molecular classification and stage classification (Figure 5A), indicating that m^6^A contributed to tumor heterogeneity. Interestingly, these different tumor subtypes have different infiltrating levels of immune cells (Figure S6), indicating m^6^A modifications shape immune responses in the tumor microenvironment and may impact cancer immunotherapy. Furthermore, the classification of these tumors was correlated with tissue origin. For example, KICH (kidney chromophobe), KIRC (kidney renal clear cell carcinoma), and KIRP (kidney renal papillary cell carcinoma), the three different types of kidney cancer, clustered in C2. Moreover, two types of lung cancer, LUAD (lung adenocarcinoma) and LUSC (lung squamous cell carcinoma), clustered in C3, and brain cancer types LGG (brain lower grade glioma) and GBM (glioblastoma multiforme) clustered in category C4 (Figure 5B). These m^6^A-regulated genes also have different clinical prognostic values. For example, patients in category C2 have a good prognosis, whereas patients in category C3 with a low infiltrating level of immune cells have a poor prognosis, indicating the influence of m^6^A on the clinical outcome of tumor patients (Figure 5C). It was proposed that m^6^A was a potential target for patients in category C3 who will not respond well to immunotherapy. Consistent with our previous conclusion, these m^6^A-regulated genes were indeed highly enriched in tumor-related pathways (Figure 5D, Figure S5C). These results suggest that m^6^A-regulated genes play different roles in diverse tumor types, resulting in distinct clinical relevance and immune status. Moreover, m^6^A may have an impact on the efficacy of immunotherapies. However, the question here is how these cancer-specific m^6^A modifications are established.

**Figure 5.**
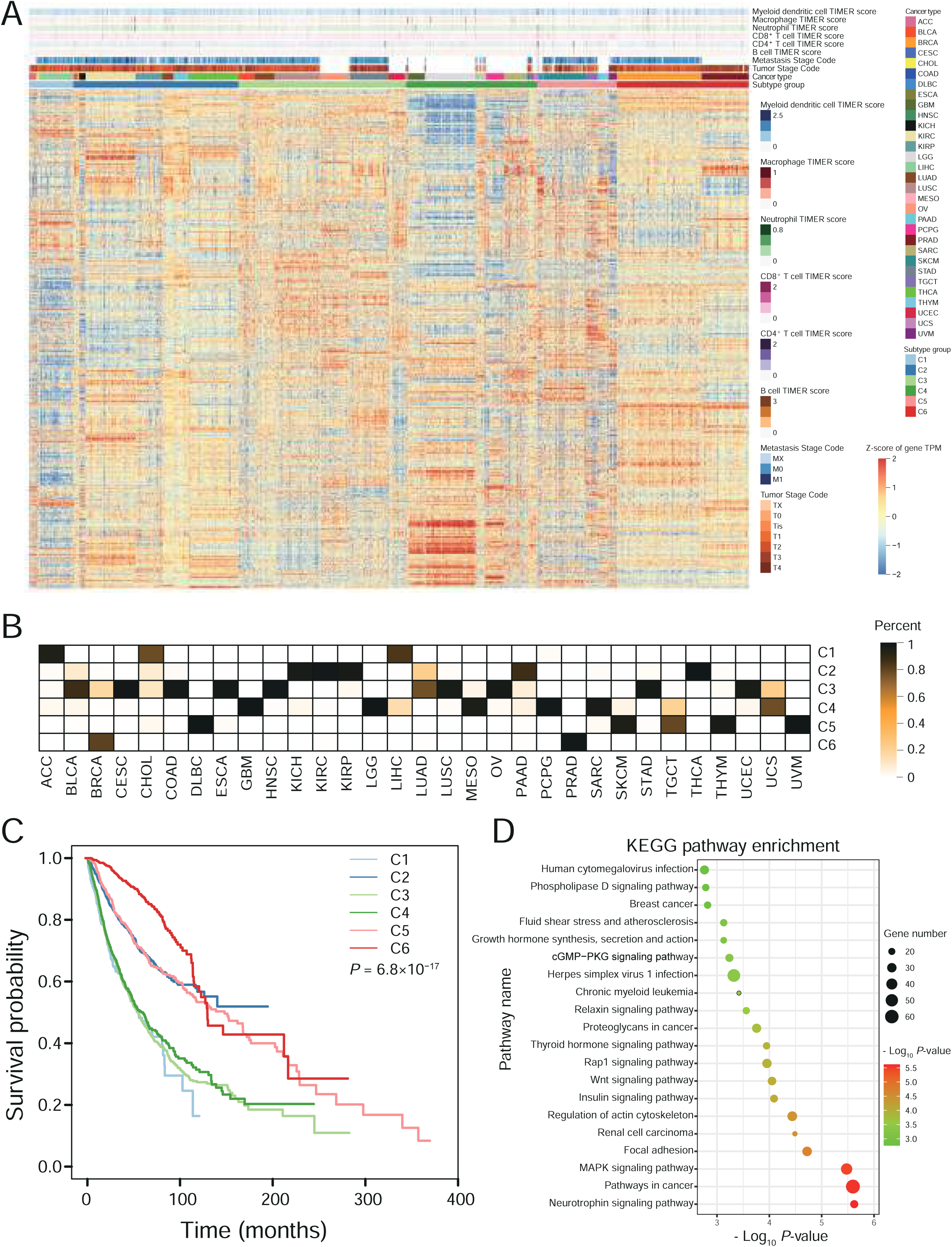
Cluster analysis of 9456 samples from 31 cancers based on the m^6^A– reg-exp genes reveals the potential of m^6^A-regulated genes in molecular subtype classification. **A**. Heatmap showing the expression of m^6^A-reg-exp genes of 31 tumors in TCGA. **B.** The proportion of each cancer in different clusters. **C.** Prognostic analysis showing different clinical outcomes of different clusters (*P* = 6.8×10^-17^). **D.** KEGG analysis of 1347 m^6^A-reg-exp genes, with bubble size representing the gene counts enriched in term and color representing the *P*-value.

### Cancer-specific m^6^A is regulated by cell-specific m^6^A regulators

Numerous studies have reported that changes in m^6^A, resulting from alterations in the expression of m^6^A regulators, play essential roles in a variety of pathological and physiological processes [21]. We previously identified 32 high-confidence cell-specific m^6^A regulators with a reasonable experimental validation rate that are responsible for global regulation and site-specific m^6^A dynamics through the interplay of classical m^6^A methyltransferase and demethylase at specific sites [21]. Herein, we calculated the Pearson correlations between the level of cancer-specific m^6^A and the expression of each m^6^A regulator. As shown in Figure 6A and 6B, the correlation coefficient values between 32 high-confidence m^6^A regulators and cancer-specific m^6^A levels were significantly higher than those of classical m^6^A regulators, indicating that cell-specific m^6^A regulators contributed more to cancer-specific m^6^A levels. We also found that cell-specific m^6^A regulators had greater CVs than classical m^6^A regulators (Figure S7). We then constructed a regulatory network based on the m^6^A regulators and cancer-type specific m^6^A levels according to correlations between the cancer-specific m^6^A levels and the expression of 32 known m^6^A regulators (Figure 6C). Among the m^6^A regulators in the regulatory network, we found that *CAPRIN1*, a novel *METTL3* co-factor [13], was positively correlated with cancer-specific m^6^A (Figure 6D). To further verify the reliability of our conclusion, we validated the regulation of *CAPRIN1* on cancer-specific m^6^A by knocking down *CAPRIN1*. We found that cancer-specific m^6^A was significantly downregulated upon *CAPRIN1* knockdown in the m^6^A-seq data (*P* = 0.006, two-tailed Wilcoxon test; Figure 6E), indicating that *CAPRIN1* regulates the installation of these cancer-specific m^6^A. We also show an example involving *TP53*, where m^6^A levels of tumor-related genes were significantly reduced in HepG2 cells with *CAPRIN1* knockdown (Figure 6F). p53 expression status is highly associated with cancer-specific survival [22], and the *CAPRIN1*-m^6^A-*TP53* axis enhances our understanding of p53-based cancer therapies. These results indicate that cancer-specific m^6^A is specifically modulated by cell-specific regulators, leading to tumor heterogeneity and influencing clinical outcomes (Figure 7).

**Figure 6.**
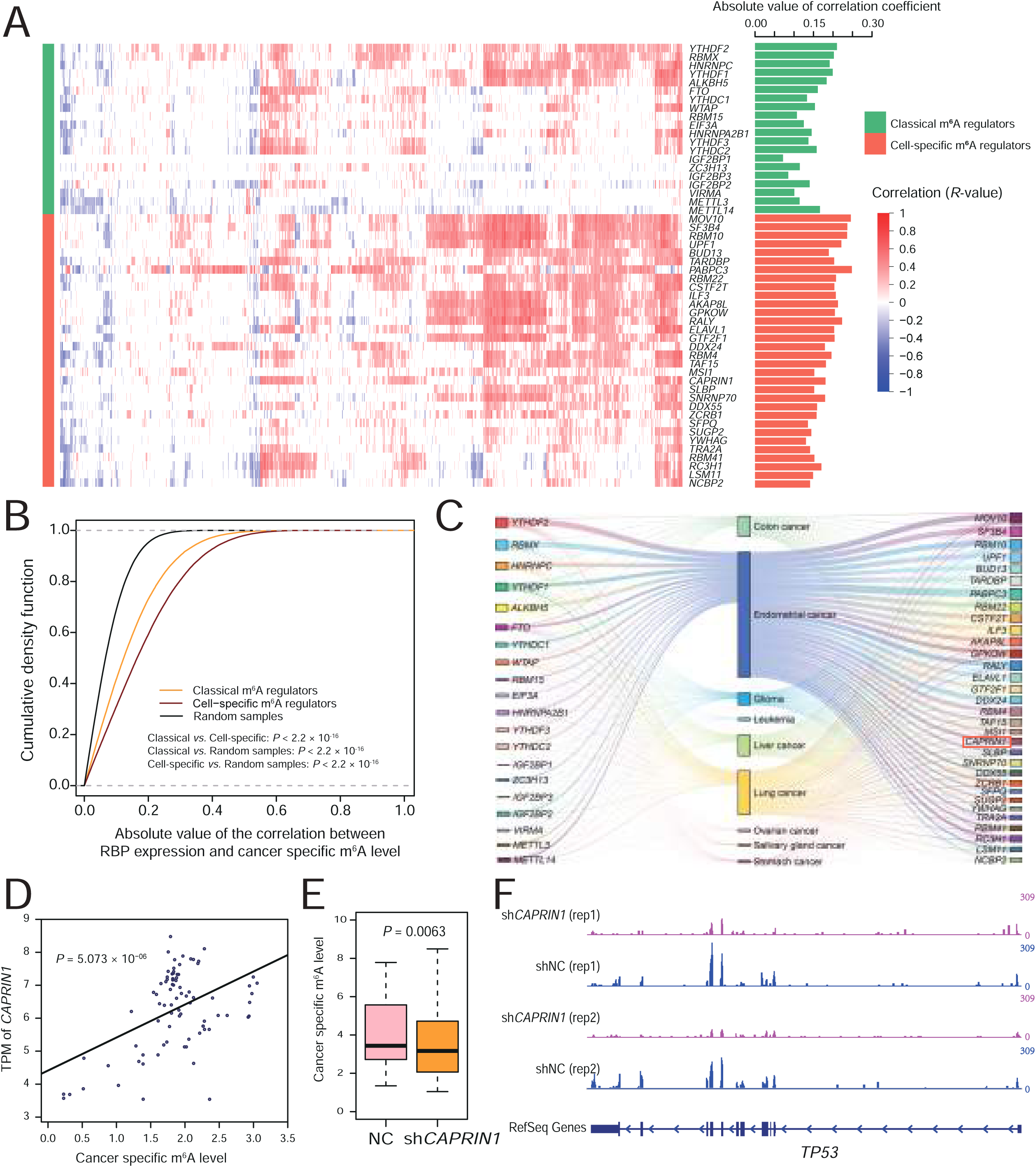
Cell-specific m^6^A regulators are involved in cancer-specific m^6^A regulation. **A**. The heatmap showing the correlation between the expression level of m^6^A regulators and corresponding cancer-specific m^6^A level, with positive correlation in red and negative correlation in blue. **B.** Plot of cumulative fraction of absolute value of the correlation coefficient between expression of two types of m^6^A regulator and corresponding cancer-specific m^6^A levels, as well as two types of m^6^A regulator and random cancer-specific m^6^A levels (*P* < 2.2 × 10^−16^). **C.** A Sankey diagram shows the network constructed based on correlation between expression level of m^6^A regulators and corresponding cancer-specific m^6^A level to identify m^6^A regulators modulating corresponding cancer specific m^6^A. **D.** Scatter plot shows the correlation between expression of the *CAPRIN1* gene and cancer-specific m^6^A level in cancer (*P* = 5.073 × 10^−6^). **E.** Boxplot showing cancer-specific m^6^A levels upon *CPARIN1* knockout (*P* = 0.0063). **F.** The track displays the read coverage of normalized IP input, highlighting the m^6^A level of the gene *TP53* in the *CPARIN1*-knockout and control group. The data range for each track is displayed on the left side (0-309).

**Figure 7.**
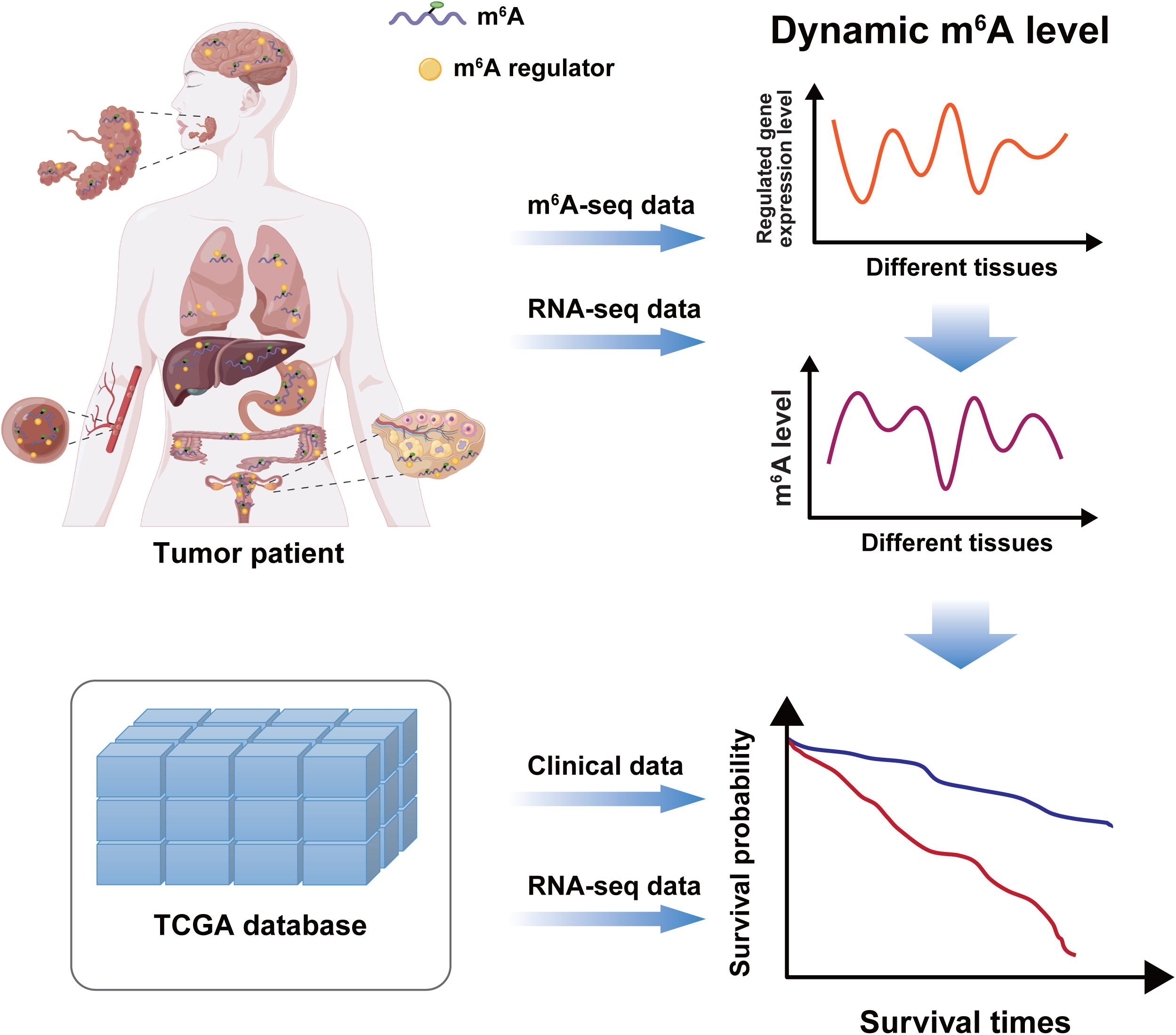
Cancer-specific m^6^A is specifically modulated by regulators, resulting in tumor heterogeneity.

## Discussion

In summary, we have demonstrated the tumor heterogeneity of m^6^A and m^6^A-reg-exp genes, which contribute to different functions and pathway enrichments. These cancer-specific m^6^A levels and functions are mainly regulated by cell-specific m^6^A regulators.

As a promising therapeutic target, m^6^A is widely involved in various biological processes in tumors, including tumorigenesis, tumor cell proliferation, apoptosis, and drug resistance [21]. For instance, *METTL3* is associated with poor prognosis in hepatocellular carcinoma (HCC) patients and promotes HCC cell proliferation through *YTHDF2*-mediated *SOCS2* transcriptional silencing [23]. *METTL14* causes the occurrence and development of leukemia by modifying *MYB/MYC*-targeted genes with m^6^A RNA, and m^6^A promotes the translation of *c-MYC*, *BCL2*, and *PTEN* genes in leukemia patients [24]. It is widely accepted that tumors exhibit heterogeneity, which influences tumor survival and response to therapy. The detailed regulatory mechanisms of m^6^A in different tumors, particularly whether the effects of m^6^A in one tumor are applicable to other tumors, remain unclear. Herein, we systematically characterized the depth and breadth of the contribution of m^6^A to interpatient tumor heterogeneity. We also systematically demonstrated downstream pan-cancer-wide m^6^A-reg-exp genes and upstream comprehensive cancer-specific m^6^A regulators in pan-cancer, promoting our understanding of the mechanisms underlying tumor heterogeneity and the role of m^6^A in tumors. We propose the following suggestions for future studies on the role of m^6^A in cancer precision therapy: (1) The functions and regulatory mechanisms of m^6^A may vary across different cancer types. (2) Small-molecule inhibition of m^6^A regulators such as *STM2457* [25], as a strategy against myeloid leukemia, may not be effective for other solid cancers. Each tumor type has its own m^6^A therapeutic targets and regulator inhibitors.

m^6^A regulators play an oncogenic role in different cancer types by targeting essential cancer-related genes [10]. A large number of studies of m^6^A regulatory mechanisms have investigated classical m^6^A regulators, such as *METTL3*, *METTL14*, *WTAP*, *METTL16*, *FTO*, *ALKBH5*, YTH family proteins, and *IGF2BPs* [26–28]; anti-cancer target drugs targeting *METTL3* and *FTO* have been proven to be effective against cancer [25, 29]. However, previous studies have shown that the m^6^A-dependent mechanism cannot be well explained by these 20 m^6^A regulators [12, 13]. In fact, we identified hundreds of novel high-confidence m^6^A regulators that were highly associated with m^6^A in different tumor types, indicating a complex regulatory system for m^6^A in tumors. This also highlighted drug targets for m^6^A in addition to the 20 m^6^A regulators. Although previous studies performed pan-cancer analysis based on the 20 classical m^6^A regulator expression profiles, several key questions remained unanswered [14, 15]. (i) Changes in gene expression levels did not fully reflect changes in m^6^A levels. (ii) m^6^A regulators such as *METTL16* function independently of m^6^A to facilitate tumorigenesis, and the effects of m^6^A in cancer may not necessarily be attributable to the effects of m^6^A regulator expression [18]. There are many more m^6^A regulators that function in cancer in addition to the 20 m^6^A regulators previously described [13, 26, 27, 30]. Our discoveries deepen the understanding of the role of m^6^A regulators based on accurate and reliable regulatory networks.

Beyond *trans-*regulation, we believe that *cis-*regulatory mechanisms also play a significant role in m^6^A. Consequently, mutations at m^6^A sites can lead to changes in m^6^A levels, which in turn affect the expression of downstream genes, further influencing the mechanism. Meng, Ren, and others have developed a series of databases that explore the interplay between m^6^A and genetic variations, such as RMVar, RMDisease and m^6^A-TSHub [31–33]. In the future, we will integrate these databases to further investigate the landscape and mechanisms of *cis-*regulation in m^6^A. Although many clinical features, especially immune dysfunction, have been associated with cancer progression, m^6^A is considered a key regulator of the immune system [34, 35]. Due to the limited availability of large samples with both m^6^A-seq and clinical data, it is challenging to investigate the reliable relationship of m^6^A with clinical features. m^6^A-express has made it possible for us to predict whole-transcriptome m^6^A-regulation of gene expression from m^6^A-seq data in TCGA. Therefore, we associated m^6^A with clinical features, including immunological characteristics, in this article. We also identified well-known transcription factors, such as *JUN* and *STAT3*, which were found to be targeted and regulated by m^6^A during tumor progression and tumor immunity [36, 37]. These factors may contribute to interpatient tumor heterogeneity and impact the effectiveness of immunotherapy, resulting in clinical challenges. Our findings regarding cell-specific m^6^A regulators modulating cancer-specific m^6^A, resulting in dysfunction of the tumor immune microenvironment, are helpful for our further understanding of cancer immunotherapy. It was proposed that immunotherapy combined with m^6^A regulator inhibitors could enhance the efficacy of immunotherapies. However, the detailed regulatory mechanism of m^6^A and the tumor immune microenvironment will require further experimental validation.

## Conclusion

In summary, our study has demonstrated the tumor heterogeneity in m^6^A and m^6^A-reg-exp genes, which contribute to different functions and pathway enrichments. These cancer-specific m^6^A levels and functions are predominantly regulated by cell-specific m^6^A regulators, resulting in tumor heterogeneity and tumor microenvironment status heterogeneity (Figure 7). Our research not only provides a landscape of the m^6^A profile in different cancer types compared to normal tissues, but also explains the clinical relevance of these specific m^6^A modifications and how these specific regulations are established. To the best of our knowledge, this is the first study based on a large number of m^6^A methylome data to propose that immunotherapy combined with m^6^A modulator inhibitors may enhance the efficacy of immunotherapy. These findings deepen our understanding of the m^6^A regulatory mechanisms in different cancer types and enhance the clinical application of m^6^A across all cancer types.

## Methods

### Data collection and processing of the m^6^A-seq data in multiple tissues

Overall, 93 raw sequence data of m^6^A-seq libraries (IP and input) were primarily downloaded from the Gene Expression Omnibus (GEO, https://www.ncbi.nlm.nih. gov/geo/) and four additional m^6^A-seq data were collected from the First Affiliated Hospital of Guangxi Medical University and deposited in SRA (PRJNA848252). Initially, a total of 97 tissue samples were collected from nine tumor types, including brain tissue, lung tissue, liver tissue, endometrium, ovarian, blood, colon, salivary gland, and stomach [38–48] (Table S1). In addition, we used the cell lines m^6^A-seq collected in previous studies [13] to verify our results, these data include seven tumor cell lines (HEC1a, HEPG2, iSLK MOLM13, MonoMac6, MT4tCELL, NB4) and three normal cell lines (MSC, NHDF, TIME) (Table S2)

We used FastQC (v0.11.9) [49] to assess the sequencing quality, and clean data were mapped to the Hg38 human reference genome by HISAT2 (v2.2.1) [50]. Then, StringTie (v1.3.3b) [51] was used for assembly and quantification of TPM (transcripts per kilobase of exon model per million mapped reads) of each annotated gene, which were then normalized by the input library. To identify accurate m^6^A sites in the nine types of tumor and adjacent normal tissues, we improved the winscore method as follows [52, 53].In detail, we performed the search for enriched m^6^A peaks by scanning each gene using sliding windows and calculating an enrichment score for each sliding window, which was modified from the method published earlier by Dominissini et al. [53]. We constructed 100 bp sliding windows with a 50 bp overlap across exon regions and determined the RPKM for each segment. Then, we designated windows with an enrichment score, or winscore, above 2 as m^6^A peaks within individual samples. To mitigate potential inaccuracies from lowly expressed windows harboring unstable winscores, we incremented each window’s RPKM by one in both IP and input datasets prior to winscore computation, thereby down-weighting windows characterized by low RPKMs. Subsequently, we amalgamated the identified m^6^A peaks across all samples for expansive analysis. We derived the m^6^A ratio of every peak by dividing the IP library’s RPKM by the input library’s RPKM. In subsequent stages of analysis, we relegated m^6^A ratios grounded on base values (the input peak’s RPKM falling below 5) as not available (NAs). Peaks designated as NAs in a majority of samples were excluded. Following this, we combined adjacent m^6^A peaks within the same gene and partitioned those extending over five continuous windows (equating 300 bp) into several peaks, each confined to a maximum of five windows.

Because different RNA interruption methods used for immunoprecipitation during preparation of different m^6^A libraries can cause changes in the short sequence signal of the same m^6^A peak, the width and center of the same m^6^A peak might be different. Therefore, we took the maximum m^6^A ratio of the combined m^6^A peaks in each sample as the final m^6^A ratio (IP/Input). Differences in activity due to different expression levels of m^6^A methylase and demethylase, as well as technical differences in immunoprecipitation efficiency, also contribute to overall m^6^A differences between samples. This dilutes and alters the signal selectively modulated by m^6^A, so we used quantile standardization, which is used to standardize the ratio of m^6^A combined with peaks in all samples [13].

### Analyses of m^6^A across cancer tissues

To compare the m^6^A peaks between cancer and normal tissues, we used the m^6^A identified in tumor tissues according to the above pipeline. To obtain the percentage of peaks enriched in representative motifs of the nine cancer tissues, HOMER software [54] was used for motif enrichment analysis, with randomly permutated sequences as the background for RNAs (HOMER parameter: line=1000, size=200). Distributions of m^6^A peaks were plotted on a mega gene with 10 bins in the 5’UTR, CDS, and 3’ UTR as previously described [13]. A radar plot was drawn using the ‘fmsb’ package implemented in R. We used bamCoverage to obtain an IP library and generate coverage tracks, with bigWig as the output. The short consecutive counting windows were set as 10 bins, and reads per kilobase per million mapped reads (RPKM) were used for normalization. With hg38 as the reference genome and *HSPD1*, *TP53*, and *JUN* as target genes, IGV (v2.8.13) was used for read coverage of *TP53* in m^6^A-seq data and *JUN* in 97 randomly selected cancer and normal samples [55]. We performed a T test (two-sided, unpaired, unequal variance) on each m^6^A site in tumor and normal tissues; 428 m^6^A sites (adjusted P < 0.05) were differentially methylated between tumor and normal tissues.

We calculated the mean and standard deviation of m^6^A modification intensity across all samples at each m^6^A site. CV is equal to the standard deviation divided by the mean value of the m^6^A sites. According to our previous report [13], sites with CV > 0.3 in specific cancer types were selected as cancer-specific m^6^A sites.

### Gene set variation analysis (GSVA) and functional annotation

To investigate the activation status of m^6^A modification patterns in different biological pathways across the nine cancer tissues, GSVA enrichment analysis was performed using the GSVA R package [56], which allows for the differential analysis of various pathways at the level of gene sets. We downloaded the gene set “c2.cp.kegg.v7.5.1 symbol” from the MSigDB database (https://www.gsea-msigdb.org/gsea/msigdb/), and an adjusted *P* < 0.05 was considered statistically significant. Functional annotation of m^6^A-related genes was performed using the ClusterProfiler R package (FDR_cutoff = 0.05).

### Identification of cancer type-specific *N^6^*-methyladenosine

We used a highly predictive and sensitive m^6^A-express computing framework based on Bayesian negative binomials [12] to evaluate the impact of m^6^A strength (IP) on the expression level (input) of each gene. With Hg38 as the reference genome, tumor and normal tissues were used for analysis of m^6^A-regulated genes (m^6^A-express parameter: DM_CUTOFF_TYPE=“pvalue”, num_ctl=2, diff_peak_pvalue=0.2, FDR=0.2, isPairedEnd=FALSE, GENE_ANNO_GTF = gtf, isGTFAnnotationFile=TRUE, DIFF_GENE_cutoff_FDR=0.2, CUTOFF_TYPE=“FDR”). Finally, 1527 m^6^A-express genes were screened by m^6^A-express. After removing duplicate genes, 1439 unique genes were considered m^6^A-regulated genes (m^6^A-reg-exp) (Table S3).

### Clustering analysis in TCGA datasets

We downloaded gene expression data and clinical data including 31 types of tumors (9456 samples), from the TCGA database. By integrating the gene expression and clinical data in TCGA (excluding LAML and READ with excessive deletion values), consensus clustering was performed to verify the effect of m^6^A-regulated genes on cancer molecular classification. Consensus clustering is an unsupervised clustering method that can distinguish samples into subtypes based on different histological datasets and allows the discovery of new disease subtypes or comparative analysis of different subtypes. To investigate the regulation of m^6^A modification on downstream gene expression, consensus clustering was performed using the “ConsensusClusterPlus” R package (k = 6) [57]. A total of 1347 m^6^A-reg-exp genes with an average TPM > 5 in TCGA were analyzed using Kyoto Encyclopedia of Genes and Genomes (KEGG) and Gene Ontology (GO) analyses. We performed functional enrichment analysis in DAVID (https://david.ncifcrf.gov/) [58], and took the top five items ranked in ascending *P*-value order as the results. GO enrichment analysis included cellular component (CC), molecular function (MF), and biological process (BP) terms.

### Sub-motif analysis

For our analysis, we first shuffled the m^6^A sub-motif sequences within all GGACA, AGACU, GGACU and GAACU m^6^A peaks for a specific sample to determine the expected number of windows containing all sub-motifs. Following this, we computed the quantity of windows that had all sub-motifs. This shuffling process was reiterated 10,000 times, yielding 10,000 expected values. To plot and compare results from different samples, we performed normalization by mean-centering the values.

## Acknowledgments

We thank Jinkai Wang from Sun Yat-sen University for the suggestions in preparing the manuscript.

## Data availability

All the data were available upon appropriate request and were deposited in public databases. Figure6 was created by Figdraw (www.figdraw.com).

## Authors’ Contributions

**Yao Lin**: Performed data collections, Conducted the analysis, Writing the manuscript, Interpreting the results, Preparing the report for publication, Revised the manuscript; **Jingyi Li**: Performed data collections, Conducted the analysis; **Shuaiyi Liang**: Revised the manuscript; **Yaxin Chen**: Partial data collection, Result collation; **Yueqi Li**: Partial data collection, Result collation; **Yixian Cun**: Partial data collection, Result collation; **Lei Tian**: Partial data collection, Result collation; **Yuanli Zhou**: Partial data collection, Result collation; **Yitong Chen**: Partial data collection, Result collation; **Jiemei Chu**: Partial data collection, Result collation; **Hubin Chen**: Partial data collection, Result collation; **Qiang Luo**: Partial data collection, Result collation; **Ruili Zheng**: Partial data collection, Result collation; **Gang Wang**: Partial data collection, Result collation; **Hao Liang**: Designed this study; **Ping Cui**: Writing the manuscript, Interpreting the results, Preparing the report for publication; **Sanqi An**: Designed this study, Performed data collections, Conducted the analysis, Writing the manuscript, Interpreting the results, Preparing the report for publication, Revised the manuscript. All authors read and approved the final manuscript.

## ORCID

ORCID 0000-0003-1728-3791 (Yao Lin)

ORCID 0009-0002-6440-5106 (Jingyi Li)

ORCID 0000-0001-8099-6159 (Shuaiyi Liang)

ORCID 0000-0002-5469-7080 (Yaxin Chen)

ORCID 0000-0002-2640-4906 (Yueqi Li)

ORCID 0000-0002-3951-942X (Yixian Cun)

ORCID 0000-0002-5268-5270 (Lei Tian)

ORCID 0000-0001-8855-489X (Yuanli Zhou)

ORCID 0000-0002-2577-2897 (Yitong Chen)

ORCID 0000-0003-2942-0490 (Jiemei Chu)

ORCID 0009-0003-0216-7138 (Hubin Chen)

ORCID 0000-0002-3753-7325 (Qiang Luo)

ORCID 0000-0002-3246-0269 (Ruili Zheng)

ORCID 0000-0001-7264-2457 (Gang Wang)

ORCID 0000-0001-7534-5124 (Hao Liang)

ORCID 0000-0001-7195-2832 (Ping Cui)

ORCID 0000-0002-3177-213X (Sanqi An)

## Figure legends

**Supplement Figure 1. m^6^A site characteristics. A.** The clustering heatmap showing the clustering effect of m^6^A-seq from different BioProject sources to calculate the abundance of m^6^A sites by improved winscore method. **B.** Comparison of overall m^6^A peak distribution between tumor (HEC1a, HEPG2, iSLK, MOLM13, MonoMac6, MT4tCELL, NB4) and normal (MSC, NHDF, TIME) cell lines. **C.** The radar map showing the percentage of classic m^6^A motifs of 7 cancer cell lines, indicated by different colored lines (Chi-Squared Test, *P* = 0.0254). **D.** The proportion of different gene types of m^6^A site in nine tumor and normal tissues. **E.** Characteristic motif of m^6^A in nine cancer tissues.

**Supplement Figure 2. Variable m^6^A sites characteristics**. **A.** The proportion of stable and variable m^6^A peaks is displayed by a stacked bar chart across all tumor and normal tissues analyzed in this study. **B.** Coefficient of variation (CV) of m^6^A level in tumor and normal tissues. The boxplot showing the CV of m^6^A level at the 5’UTR (*, *P* < 0.05; **, *P* < 0.01; ****, *P* < 0.0001). **C.** The track shows the m^6^A coverage of the *HSPD1* gene from randomly selected samples among 97 cancer and normal subjects. The data range for each track is displayed on the left side (0-96). **D.** Differences in m^6^A between cancer and normal samples (*P* < 2.2 × 10^−16^).

**Supplement Figure 3. GSVA enrichment analysis revealed activation status of different cancer-specific m^6^A-related KEGG pathways**. After being Z-score normalized, the GSVA sample-wise gene set enrichment scores are used to plot a heatmap.

**Supplement Figure 4. GO enrichment analysis of cancer type-specific m^6^A across nine cancer types**. The x-axis represents the significance level of pathway enrichment. A. Cellular component. B. molecular function.

**Supplement Figure 5. Classification of 31 cancer types based on m^6^A-reg-exp gene profiles**. **A.** Consensus clustering matrix of m^6^A-reg-exp genes in 31 tumors in TCGA for k = 6. **B.** Consensus clustering CDF for k = 2 to k = 12. **C.** Biological process of 1347 m^6^A-reg-exp genes by GO enrichment analysis (Biological Progress).

**Supplement Figure 6. Plot of cumulative fraction and boxplot in 6 subtypes of immune cells A.** B cell TIMER score, **B.** CD4+ T cell TIMER score, **C.** CD8+ T cell TIMER score, **D.** Macrophage TIMER score, **E.** Myeloid dendritic cell TIMER score, **F.** Neutrophil TIMER score calculated by TIMER.

Supplement Figure 7. Heatmap of expression of classical m^6^A regulators and cell specific m^6^A regulators in 31 tumor samples in TCGA. Bar graph shows the coefficient of variation of corresponding genes.

Supplement Table 1. Data collection of 9 types of cancer and normal tissues m^6^A-seq.

Supplement Table 2. Data collection of 7 types of cancer cell lines m^6^A-seq and 3 types of normal cell lines m^6^A-seq.

Supplement Table 3. m^6^A-express result delineates cancer specific genes that are regulated by m^6^A in distinct cancer types, alongside their expression levels across diverse samples.

## Reference

[1] ICGC/TCGA Pan-Cancer Analysis of Whole Genomes Consortium. Pan-cancer analysis of whole genomes. Nature 2020; (578):82–93.

[2] Chiu HS, Somvanshi S, Patel E, Chen TW, Singh VP, Zorman B, et al. Pan-Cancer Analysis of lncRNA Regulation Supports Their Targeting of Cancer Genes in Each Tumor Context. Cell Rep 2018; (23):297–312 e12.

[3] Chen H, Li C, Peng X, Zhou Z, Weinstein JN, Cancer Genome Atlas Research N, et al. A Pan-Cancer Analysis of Enhancer Expression in Nearly 9000 Patient Samples. Cell 2018; (173):386–99 e12.

[4] Bin Lim S, Chua MLK, Yeong JPS, Tan SJ, Lim WT, Lim CT. Pan-cancer analysis connects tumor matrisome to immune response. NPJ Precis Oncol 2019; (3):15.

[5] Cheng M, Sheng L, Gao Q, Xiong Q, Zhang H, Wu M, et al. The m(6)A methyltransferase METTL3 promotes bladder cancer progression via AFF4/NF-kappaB/MYC signaling network. Oncogene 2019; (38):3667–80.

[6] Chen Y, Zhou C, Sun Y, He X, Xue D. m(6)A RNA modification modulates gene expression and cancer-related pathways in clear cell renal cell carcinoma. Epigenomics 2020; (12):87–99.

[7] Schumann U, Shafik A, Preiss T. METTL3 Gains R/W Access to the Epitranscriptome. Molecular Cell 2016; (62):323–4.

[8] Huang H, Weng H, Sun W, Qin X, Shi H, Wu H, et al. Recognition of RNA N(6)-methyladenosine by IGF2BP proteins enhances mRNA stability and translation. Nature Cell Biology 2018; (20):285–95.

[9] Zhang C, Huang S, Zhuang H, Ruan S, Zhou Z, Huang K, et al. YTHDF2 promotes the liver cancer stem cell phenotype and cancer metastasis by regulating OCT4 expression via m6A RNA methylation. Oncogene 2020; (39):4507–18.

[10] Barbieri I, Kouzarides T. Role of RNA modifications in cancer. Nat Rev Cancer 2020; (20):303–22.

[11] Nombela P, Miguel-Lopez B, Blanco S. The role of m(6)A, m(5)C and Psi RNA modifications in cancer: Novel therapeutic opportunities. Mol Cancer 2021; (20):18.

[12] Zhang T, Zhang SW, Zhang SY, Gao SJ, Chen Y, Huang Y. m6A-express: uncovering complex and condition-specific m6A regulation of gene expression. Nucleic Acids Res 2021; (49):e116.

[13] An S, Huang W, Huang X, Cun Y, Cheng W, Sun X, et al. Integrative network analysis identifies cell-specific trans regulators of m6A. Nucleic Acids Res 2020; (48):1715–29.

[14] Shen S, Zhang R, Jiang Y, Li Y, Lin L, Liu Z, et al. Comprehensive analyses of m6A regulators and interactive coding and non-coding RNAs across 32 cancer types. Mol Cancer 2021; (20):67.

[15] Li Y, Xiao J, Bai J, Tian Y, Qu Y, Chen X, et al. Molecular characterization and clinical relevance of m(6)A regulators across 33 cancer types. Mol Cancer 2019; (18):137.

[16] Larivera S, Meister G. Domain confusion 2: m(6)A-independent role of YTHDC2. Molecular Cell 2022; (82):1608–9.

[17] Liu P, Li F, Lin J, Fukumoto T, Nacarelli T, Hao X, et al. m(6)A-independent genome-wide METTL3 and METTL14 redistribution drives the senescence-associated secretory phenotype. Nature Cell Biology 2021; (23):355–65.

[18] Su R, Dong L, Li Y, Gao M, He PC, Liu W, et al. METTL16 exerts an m(6)A-independent function to facilitate translation and tumorigenesis. Nature Cell Biology 2022; (24):205–16.

[19] Dominissini D, Moshitch-Moshkovitz S, Salmon-Divon M, Amariglio N, Rechavi G. Transcriptome-wide mapping of N(6)-methyladenosine by m(6)A-seq based on immunocapturing and massively parallel sequencing. Nature Protocols 2013; (8):176–89.

[20] Garcia-Campos MA, Edelheit S, Toth U, Safra M, Shachar R, Viukov S, et al. Deciphering the “m(6)A Code” via Antibody-Independent Quantitative Profiling. Cell 2019; (178):731–47 e16.

[21] Jiang X, Liu B, Nie Z, Duan L, Xiong Q, Jin Z, et al. The role of m6A modification in the biological functions and diseases. Signal Transduct Target Ther 2021; (6):74.

[22] Oh HJ, Bae JM, Wen X, Jung S, Kim Y, Kim KJ, et al. p53 expression status is associated with cancer-specific survival in stage III and high-risk stage II colorectal cancer patients treated with oxaliplatin-based adjuvant chemotherapy. British Journal of Cancer 2019; (120):797–805.

[23] Chen M, Wei L, Law CT, Tsang FH, Shen J, Cheng CL, et al. RNA N6-methyladenosine methyltransferase-like 3 promotes liver cancer progression through YTHDF2-dependent posttranscriptional silencing of SOCS2. Hepatology 2018; (67):2254–70.

[24] Liu X, Du Y, Huang Z, Qin H, Chen J, Zhao Y. Insights into roles of METTL14 in tumors. Cell Proliferation 2022; (55):e13168.

[25] Yankova E, Blackaby W, Albertella M, Rak J, De Braekeleer E, Tsagkogeorga G, et al. Small-molecule inhibition of METTL3 as a strategy against myeloid leukaemia. Nature 2021; (593):597–601.

[26] Liu M, Xu K, Saaoud F, Shao Y, Zhang R, Lu Y, et al. 29 m(6)A-RNA Methylation (Epitranscriptomic) Regulators Are Regulated in 41 Diseases including Atherosclerosis and Tumors Potentially via ROS Regulation – 102 Transcriptomic Dataset Analyses. J Immunol Res 2022; (2022):1433323.

[27] Zhu J, Xiao J, Wang M, Hu D. Pan-Cancer Molecular Characterization of m(6)A Regulators and Immunogenomic Perspective on the Tumor Microenvironment. Front Oncol 2020; (10):618374.

[28] Zhang Y, Hamada M. Identification of m(6)A-Associated RNA Binding Proteins Using an Integrative Computational Framework. Front Genet 2021; (12):625797.

[29] Su R, Dong L, Li Y, Gao M, Han L, Wunderlich M, et al. Targeting FTO Suppresses Cancer Stem Cell Maintenance and Immune Evasion. Cancer Cell 2020; (38):79–96 e11.

[30] Xu T, Wu X, Wong CE, Fan S, Zhang Y, Zhang S, et al. FIONA1-Mediated m(6) A Modification Regulates the Floral Transition in Arabidopsis. Adv Sci (Weinh) 2022; (9):e2103628.

[31] Luo X, Li H, Liang J, Zhao Q, Xie Y, Ren J, et al. RMVar: an updated database of functional variants involved in RNA modifications. Nucleic Acids Res 2021; (49):D1405–D12.

[32] Song B, Wang X, Liang Z, Ma J, Huang D, Wang Y, et al. RMDisease V2.0: an updated database of genetic variants that affect RNA modifications with disease and trait implication. Nucleic Acids Res 2023; (51):D1388–D96.

[33] Song B, Huang D, Zhang Y, Wei Z, Su J, Pedro de Magalhaes J, et al. m6A-TSHub: Unveiling the Context-specific m(6)A Methylation and m6A-affecting Mutations in 23 Human Tissues. Genomics Proteomics & Bioinformatics 2022.

[34] Chong W, Shang L, Liu J, Fang Z, Du F, Wu H, et al. m(6)A regulator-based methylation modification patterns characterized by distinct tumor microenvironment immune profiles in colon cancer. Theranostics 2021; (11):2201–17.

[35] Shulman Z, Stern-Ginossar N. The RNA modification N(6)-methyladenosine as a novel regulator of the immune system. Nat Immunol 2020; (21):501–12.

[36] Zhu W, Wang JZ, Wei JF, Lu C. Role of m6A methyltransferase component VIRMA in multiple human cancers (Review). Cancer Cell Int 2021; (21):172.

[37] Yang Z, Cai Z, Yang C, Luo Z, Bao X. ALKBH5 regulates STAT3 activity to affect the proliferation and tumorigenicity of osteosarcoma via an m6A-YTHDF2-dependent manner. EBioMedicine 2022; (80):104019.

[38] Hou J, Zhang H, Liu J, Zhao Z, Wang J, Lu Z, et al. YTHDF2 reduction fuels inflammation and vascular abnormalization in hepatocellular carcinoma. Mol Cancer 2019; (18):163.

[39] An S, Xie Z, Liao Y, Jiang J, Dong W, Yin F, et al. Systematic analysis of clinical relevance and molecular characterization of m(6)A in COVID-19 patients. Genes Dis 2022; (9):1170–3.

[40] Zhang Z, Zhan Q, Eckert M, Zhu A, Chryplewicz A, De Jesus DF, et al. RADAR: differential analysis of MeRIP-seq data with a random effect model. Genome Biol 2019; (20):294.

[41] Choe J, Lin S, Zhang W, Liu Q, Wang L, Ramirez-Moya J, et al. mRNA circularization by METTL3-eIF3h enhances translation and promotes oncogenesis. Nature 2018; (561):556–60.

[42] Liu J, Eckert MA, Harada BT, Liu SM, Lu Z, Yu K, et al. m(6)A mRNA methylation regulates AKT activity to promote the proliferation and tumorigenicity of endometrial cancer. Nat Cell Biol 2018; (20):1074–83.

[43] Niu X, Xu J, Liu J, Chen L, Qiao X, Zhong M. Landscape of N(6)-Methyladenosine Modification Patterns in Human Ameloblastoma. Front Oncol 2020; (10):556497.

[44] Liu J, Li K, Cai J, Zhang M, Zhang X, Xiong X, et al. Landscape and Regulation of m(6)A and m(6)Am Methylome across Human and Mouse Tissues. Molecular Cell 2020; (77):426–40 e6.

[45] Han Z, Ren H, Sun J, Jin L, Wang Q, Guo C, et al. Integrated weighted gene coexpression network analysis identifies Frizzled 2 (FZD2) as a key gene in invasive malignant pleomorphic adenoma. J Transl Med 2022; (20):15.

[46] Li Z, Weng H, Su R, Weng X, Zuo Z, Li C, et al. FTO Plays an Oncogenic Role in Acute Myeloid Leukemia as a N(6)-Methyladenosine RNA Demethylase. Cancer Cell 2017; (31):127–41.

[47] Zhou Y, Zhou H, Shi J, Guan A, Zhu Y, Hou Z, et al. Decreased m6A Modification of CD34/CD276(B7-H3) Leads to Immune Escape in Colon Cancer. Front Cell Dev Biol 2021; (9):715674.

[48] Han Z, Yang B, Wang Q, Hu Y, Wu Y, Tian Z. Comprehensive analysis of the transcriptome-wide m(6)A methylome in invasive malignant pleomorphic adenoma. Cancer Cell Int 2021; (21):142.

[49] S A. FastQC: a quality control tool for high throughput sequence data http://www.bioinformatics.babraham.ac.uk/projects/fastqc. 2010.

[50] Kim D, Langmead B, Salzberg SL. HISAT: a fast spliced aligner with low memory requirements. Nature Methods 2015; (12):357–60.

[51] Pertea M, Pertea GM, Antonescu CM, Chang TC, Mendell JT, Salzberg SL. StringTie enables improved reconstruction of a transcriptome from RNA-seq reads. Nature Biotechnology 2015; (33):290–5.

[52] Batista PJ, Molinie B, Wang J, Qu K, Zhang J, Li L, et al. m(6)A RNA modification controls cell fate transition in mammalian embryonic stem cells. Cell Stem Cell 2014; (15):707–19.

[53] Dominissini D, Moshitch-Moshkovitz S, Schwartz S, Salmon-Divon M, Ungar L, Osenberg S, et al. Topology of the human and mouse m6A RNA methylomes revealed by m6A-seq. Nature 2012; (485):201–6.

[54] Heinz S, Benner C, Spann N, Bertolino E, Lin YC, Laslo P, et al. Simple combinations of lineage-determining transcription factors prime cis-regulatory elements required for macrophage and B cell identities. Molecular Cell 2010; (38):576–89.

[55] Thorvaldsdottir H, Robinson JT, Mesirov JP. Integrative Genomics Viewer (IGV): high-performance genomics data visualization and exploration. Brief Bioinform 2013; (14):178–92.

[56] Hanzelmann S, Castelo R, Guinney J. GSVA: gene set variation analysis for microarray and RNA-seq data. BMC Bioinformatics 2013; (14):7.

[57] Wilkerson MD, Hayes DN. ConsensusClusterPlus: a class discovery tool with confidence assessments and item tracking. Bioinformatics 2010; (26):1572–3.

[58] Huang DW, Sherman BT, Tan Q, Kir J, Liu D, Bryant D, et al. DAVID Bioinformatics Resources: expanded annotation database and novel algorithms to better extract biology from large gene lists. Nucleic Acids Res 2007; (35):W169–75.

