## Supplementary figures and images for "Pan-cancer Analysis Reveals m^6^A Variation and Cell-specific Regulatory Network in Different Cancer Types"

### Supplementary Figure 1

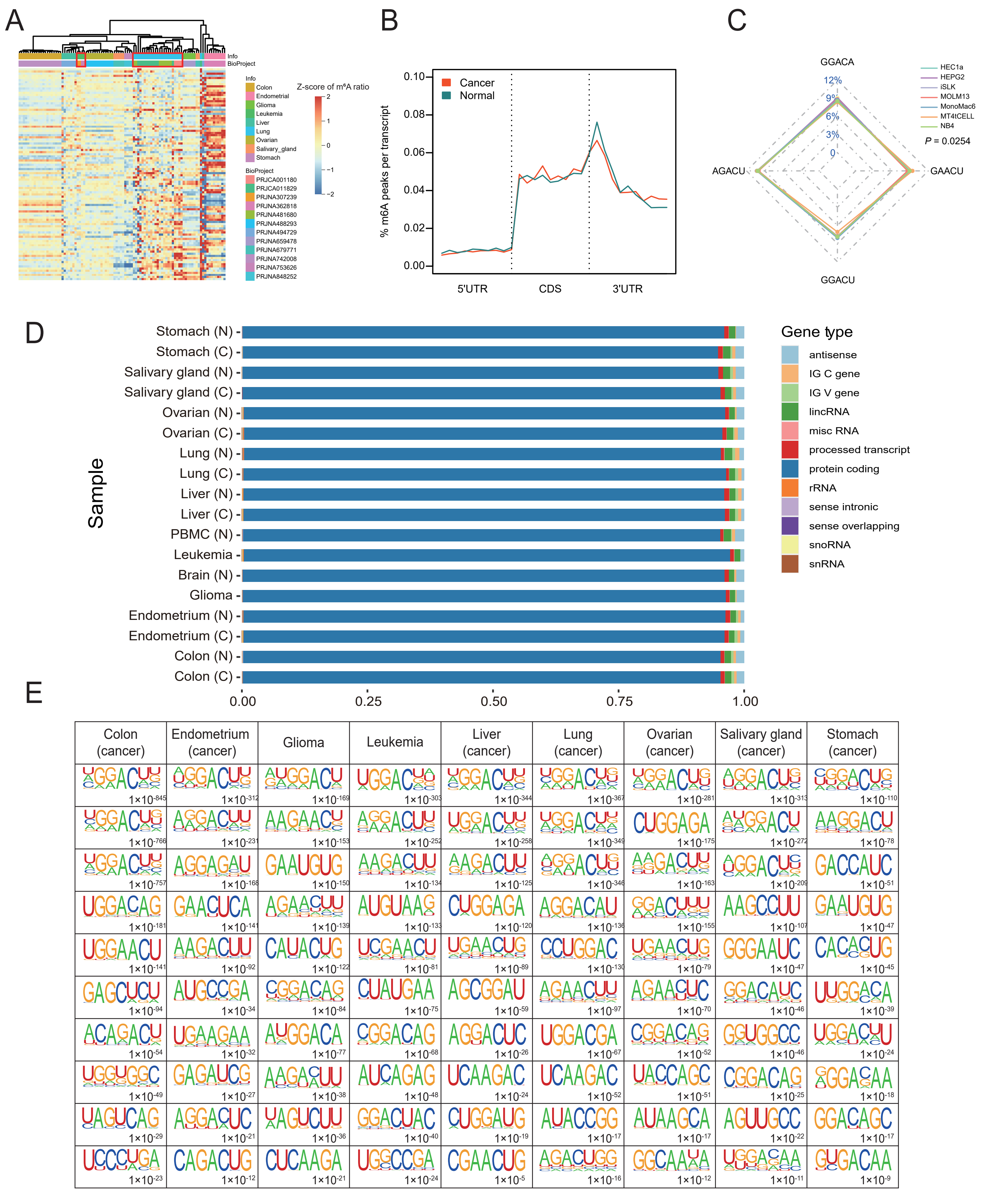

### Supplementary Figure 2

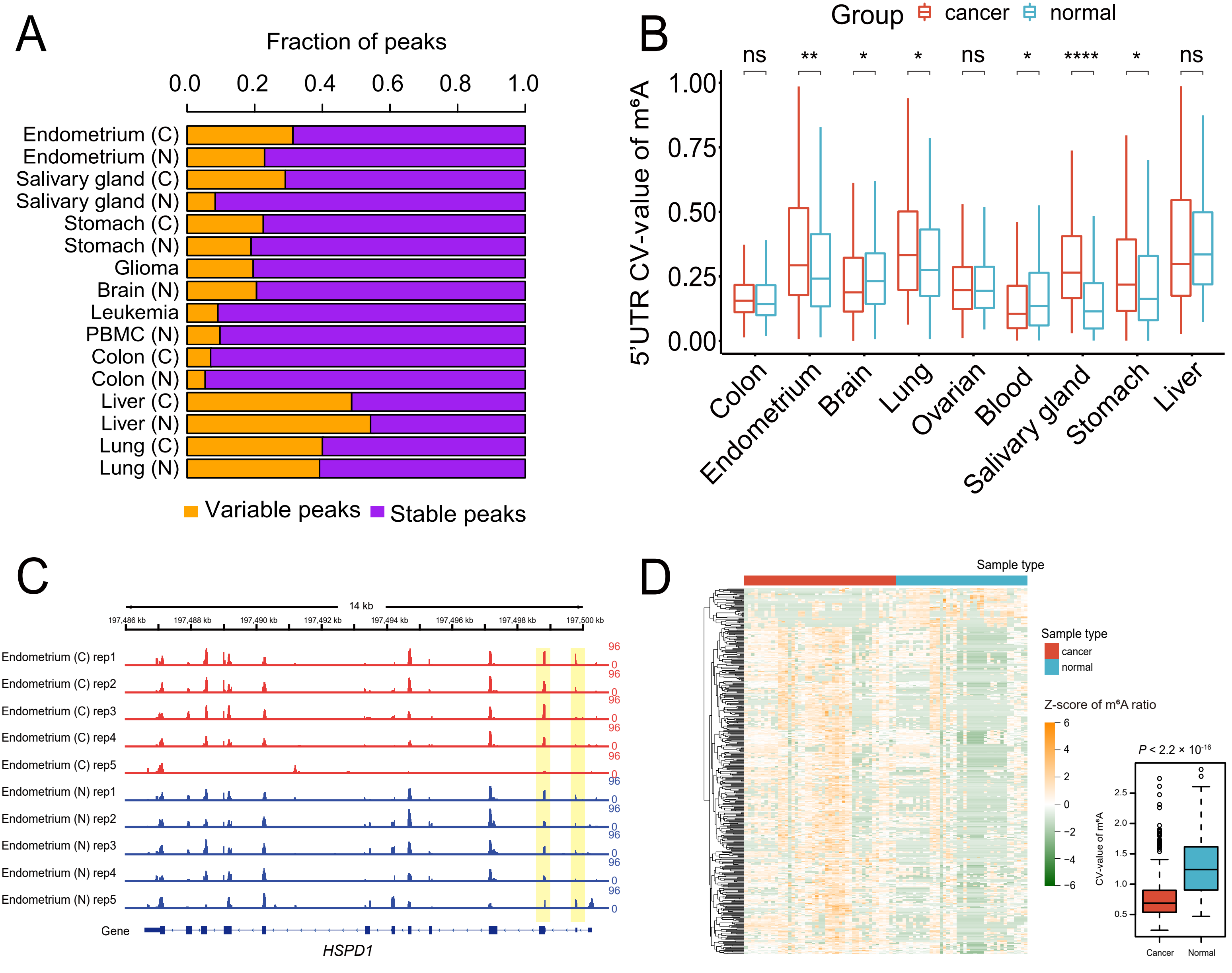

### Supplementary Figure 3

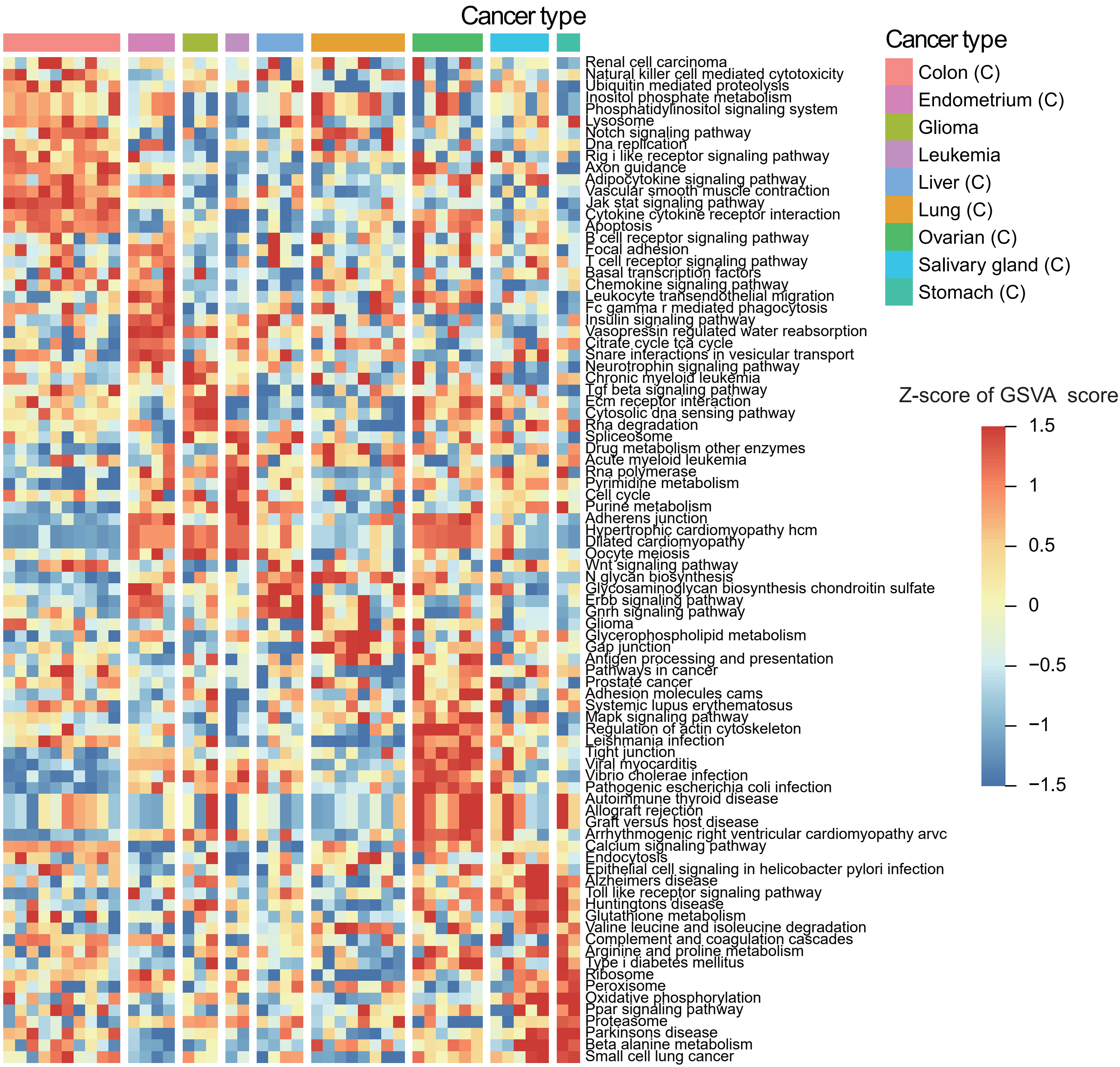

### Supplementary Figure 4

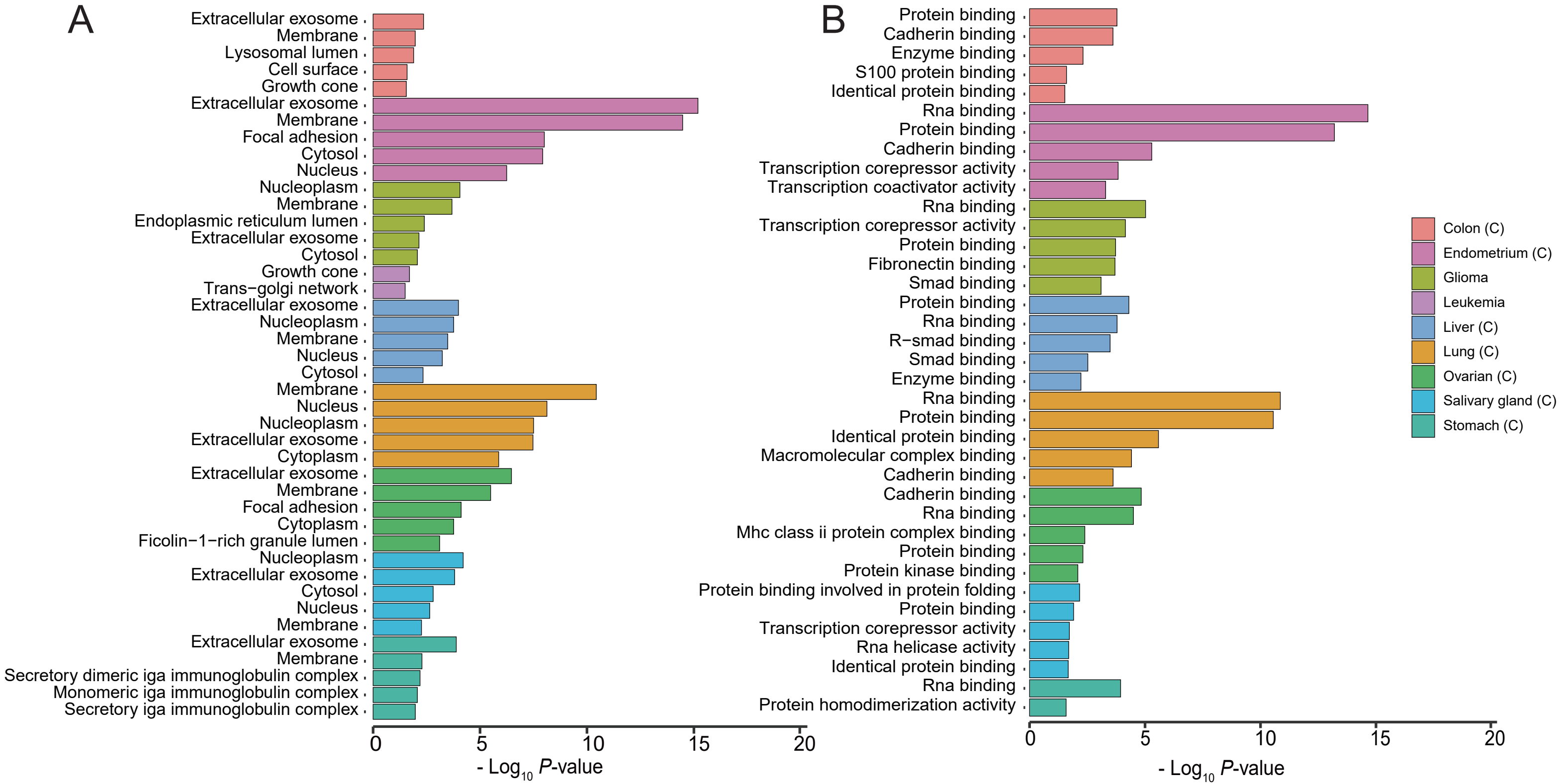

### Supplementary Figure 5

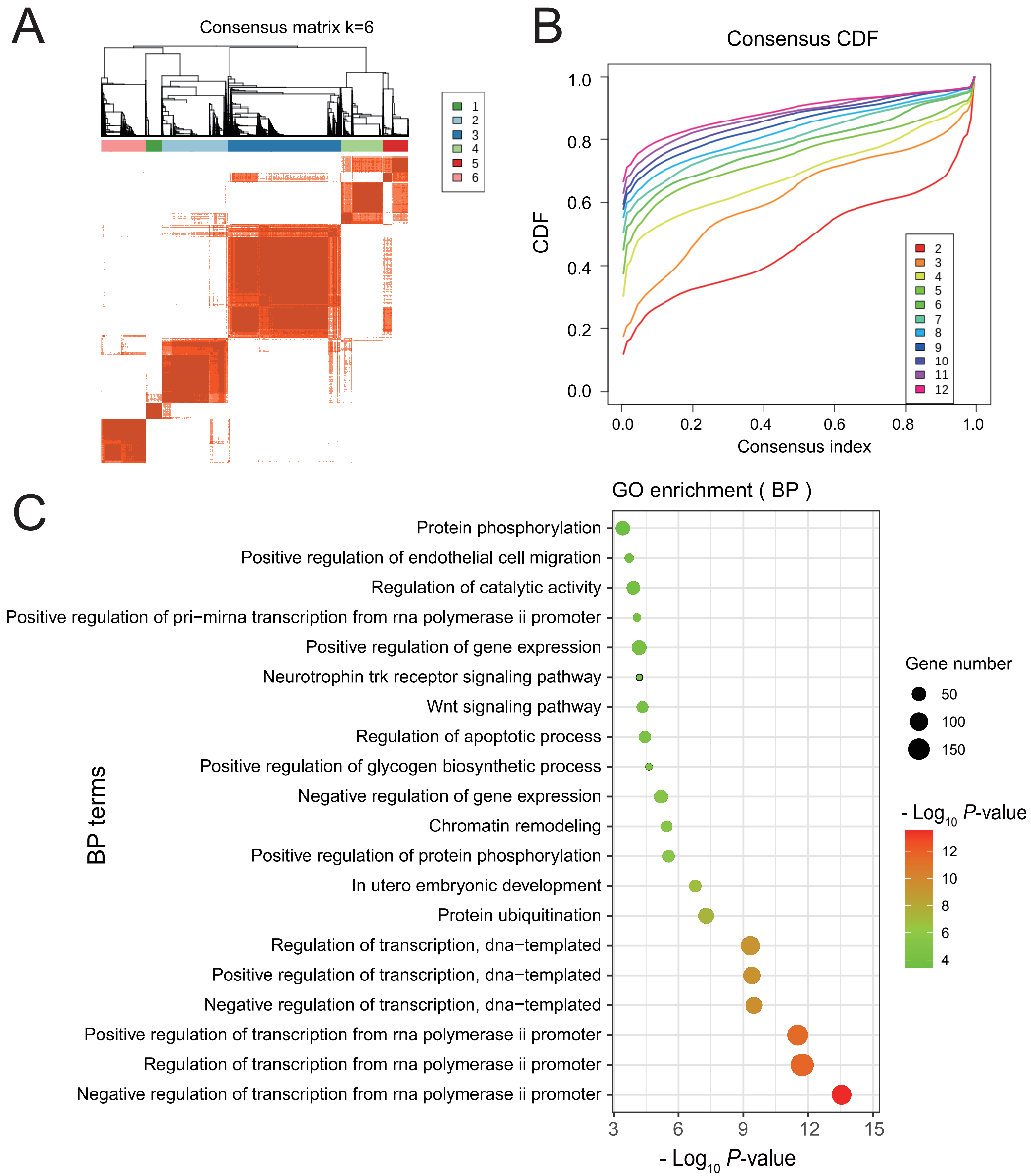

### Supplementary Figure 6

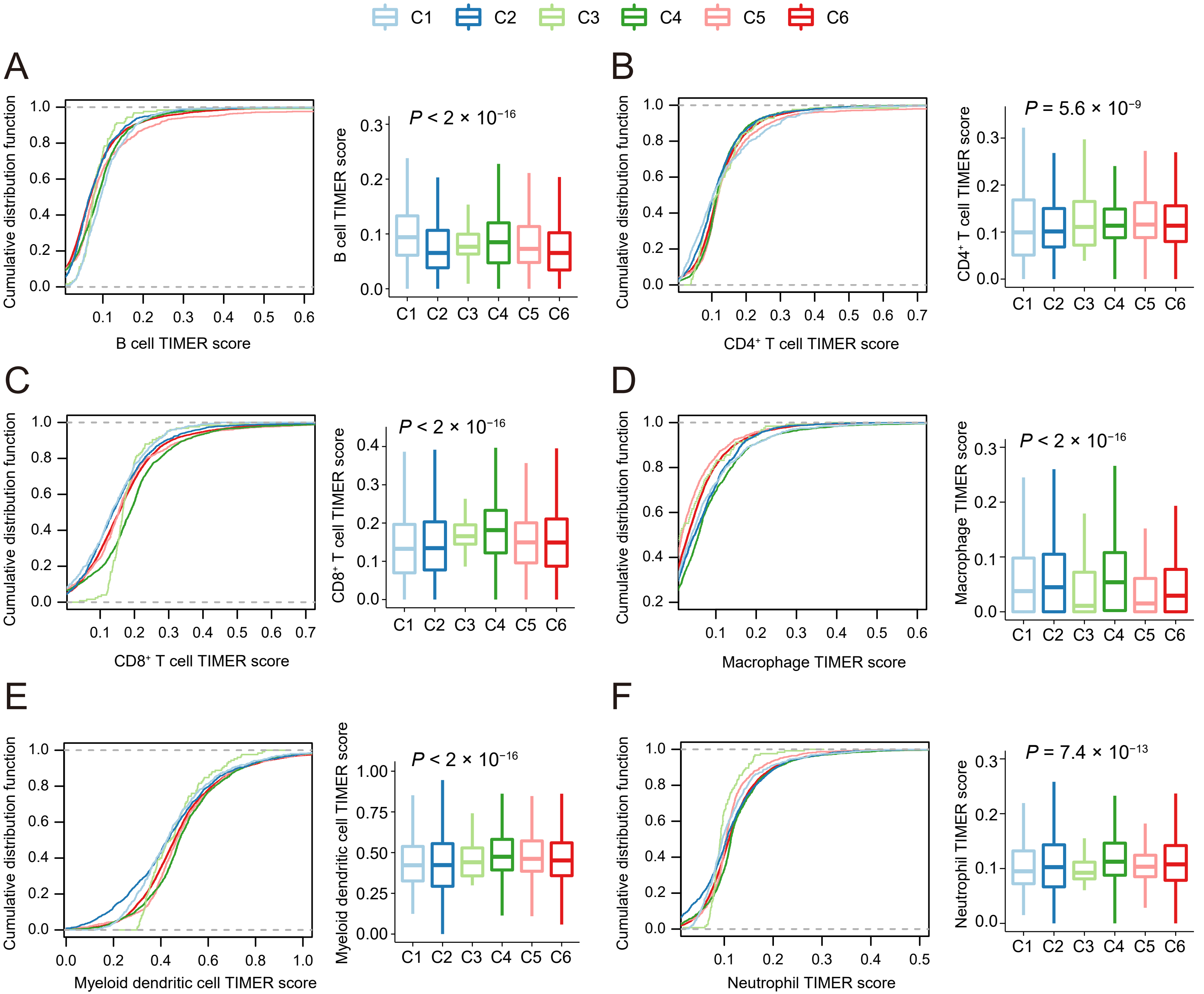

### Supplementary Figure 7

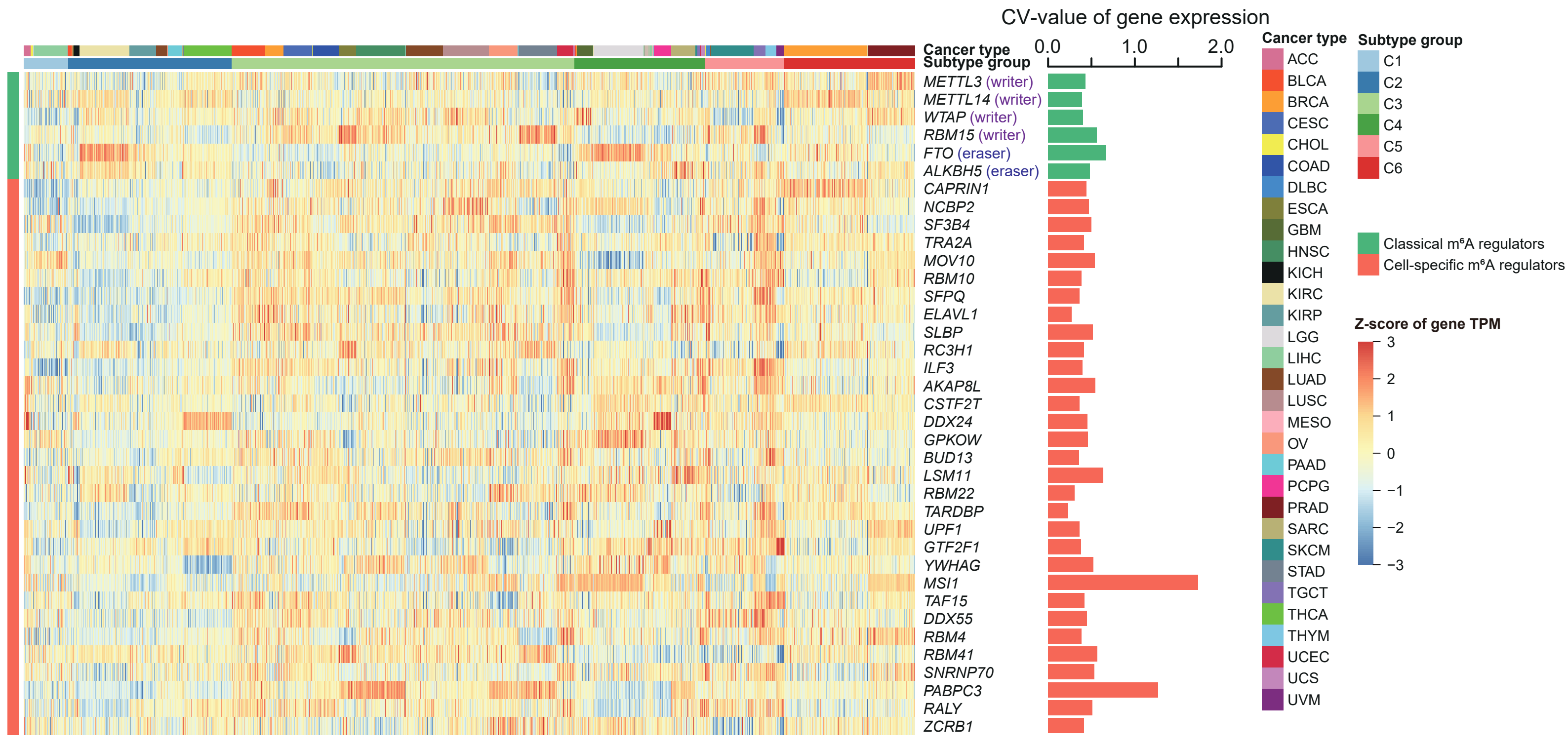
